# Strengthened mutualistic adaptation between teosinte and its rhizosphere biota in cold climates

**DOI:** 10.1101/2021.04.20.440703

**Authors:** Anna M. O’Brien, Ruairidh J.H. Sawers, Jaime Gasca-Pineda, Ivan Baxter, Luis E. Eguiarte, Jeffrey Ross-Ibarra, Sharon Y. Strauss

**Author notes:** Address correspondence to Anna M. O’Brien.

## Abstract

- While abiotic environments consistently shape local adaptation, the strength of local adaptation to biotic interactions may vary more. One theory, COCO (CO-evolutionary Outcomes across Conditionality), predicts it may be strongest where species experience greater stress, because stress increases fitness impacts of species interactions. For example, in plant interactions with rhizosphere biota, positive outcomes increase with stress from low soil fertility, drought and cold.
- To investigate the influence of abiotic stress gradients on adaptation between plants and rhizosphere biota, we used a greenhouse common garden experiment recombining teosinte, *Zea mays* ssp. *mexicana* (wild relative of maize), and rhizosphere biota, collected across a stress gradient (elevational variation in temperature, precipitation, and nutrients).
- We found stronger local adaptation between teosinte and rhizosphere biota from colder, more stressful sites, as expected by COCO. However, biota from less stressful, warmer sites provided greater average benefits across teosinte populations. Links between plant traits and 20-element profiles of plant leaves explained fitness variation, persisted in the field, were influenced by both plants and biota, and largely reflected patterns of local adaptation.
- In sum, we uncovered greater local adaptation to biotic interactions in colder sites, and that both plants and rhizosphere biota affect the expression of plant phenotypes.

## Introduction

The striking power of both abiotic and biotic selective forces in evolution has been well-documented, yet meta-analyses reveal that while abiotic forces consistently drive strong local adaptation to sites across species and systems, local adaption to biotic interactions is inconsistent in strength (Briscoe Runquist et al., 2020; Hargreaves et al., 2020). This is not unexpected: fitness impacts of biotic interactions vary across abiotic conditions as the impacts of individual interactions or the composition of suites of interacting species shift (Cushman and Whitham, 1989; Strauss and Irwin, 2004; Chamberlain et al., 2014; Trøjelsgaard et al., 2015; Kemp et al., 2017), driving mosaics of co-evolution and co-adaptation (Thompson, 1982, 2005). One well-known pattern of shifting outcomes is the strengthening of benefits between partners across gradients of abiotic stress, heightening mutualisms (Johnson, 1993; Schwartz and Hoeksema, 1998; Bever, 2015) and attenuating competition or shifting it to facilitation (Bertness and Callaway, 1994; Callaway et al., 2002; He and Bertness, 2014). Altered adaptation to species interactions is then expected to follow such shifts in outcomes (Bronstein, 2009; O’Brien et al., 2018). Specifically, the “(co)evolutionary outcomes of conditionaliy” (COCO) hypothesis predicts increased mutualism and local adaptation in one or both interacting species in stressful sites where the interaction ameliorates the stressor’s effect (O’Brien et al., 2018).

For plants, one important class of biotic interaction is with the diverse community of organisms that live in, on, or near their roots (Hiltner, 1904; Bais et al., 2006; Raaijmakers et al., 2009; Lundberg et al., 2012; Toju et al., 2014). Though these interactions are primarily mutualisms involving exchange of plant photosynthetically-fixed carbon for nutritional benefits from biota (Smith and Read, 2008), outcomes for plants may sometimes be costly (e.g. Berg and Smalla, 2009; Smith and Read, 2008; Anacker et al., 2014). Like patterns across species, local adaptation between plants and components of their rhizosphere biota is variable in strength (Rúa et al., 2016). Because plant-rhizosphere interactions provide positive outcomes through ameliorating abiotic stress, they may shift to negative outcomes in the absence of that stress (Johnson, 1993), leading to the prediction of COCO and other theoretical frameworks that there should be stronger mutualistic adaptation between plants and microbes in high-stress sites such as those lacking in soil nutrients (Bever, 2015; O’Brien et al., 2018). In one tantalizing example, mycorrhizal fungi positively affect plants by alleviating phosphorus stress, and greater local adaptation between plants and fungi was observed in phosphorus-deficient sites compared to sites with less phosphorus stress (Johnson et al., 2010).

Here, we address the influence of abiotic environments on local adaptation between teosinte, *Zea mays* ssp. *mexicana*, a wild relative of domesticated maize (*Zea mays* ssp. *mays*) from the highlands of central Mexico (Sánchez and Corral, 1997) and its rhizosphere biota. We experimentally combined teosinte plants and rhizosphere biota from sites spanning an elevational range that also captured gradients in soil fertility, temperature, and precipitation (O’Brien et al., 2019). These gradients may have synergistic effects: cold stress in plants is physiologically driven by water and nutrients as roots function poorly in the cold, leading to nutrient deficiencies and wilting (Bloom et al., 2004; Zhu et al., 2009), potentially exacerbating effects of dry or nutrient-poor sites. Rhizosphere biota can alleviate drought (Kivlin et al., 2013), cold (Zhu et al., 2009) and nutrient stress (Smith et al., 2010), and may therefore be most beneficial in dry, nutrient-poor, and cold sites. COCO predicts the most evolved mutualistic benefits where interactions have the most beneficial outcomes. Benefits can be general and provided to any interacting partner, or they may be locally adapted and provided only to partners from the local population (O’Brien et al., 2018). We hypothesized that *1)* biota from the most stressful sites (cold, dry, nutrient-poor) would provide the most general and locally adapted benefits.

In teosinte, many of the traits that underlie adaptive differentiation across elevation or cold stress (including phenology and height, Hufford et al., 2013; O’Brien et al., 2019; Fustier et al., 2019) also shift in response to changes in rhizosphere communities (O’Brien et al., 2019). Plant-rhizosphere interactions may simultaneously influence many different elements in plant tissues (e.g. the ionome, a 20-element profile, Baxter et al., 2008; Ramírez-Flores et al., 2017), and microbially driven shifts in plant tissue element concentrations are linked to shifts in plant traits from root architecture to flowering time (Desbrosses and Stougaard, 2011; Bulgarelli et al., 2013; Paszkowski and Gutjahr, 2013; Lu et al., 2018), suggesting an interplay between rhizosphere nutrient provisioning, and the expression of adaptive phenotypes. To investigate any such interplay between teosinte and rhizosphere biota, we identified plant phenotypes that co-varied with elemental profiles. We hypothesized that *2)* adaptation associated with stress gradients (cold, fertility, precipitation) would shape co-varying plant traits and element profiles, i.e. that patterns in traits and elements would reflect general and locally adapted benefits from biota.

Finally, benefits provided to plants by biota should be greater when conditions match the local environment from which plants and biota were sourced and to which they may be locally adapted (Johnson et al., 2010; Lau and Lennon, 2012). We therefore measured element profiles and a subset of traits in the field, and tested whether *3)* rhizosphere biota both provide greater benefits to teosinte at more stressful sites (cold, dry, nutrient-poor), and shift traits and elemental profiles as observed in the greenhouse.

## Materials and Methods

### Characterization of field sites and collections

We selected 10 populations of teosinte from central Mexico across its elevational range (Figure S1) that we expected to differ in soil fertility (based on underlying geology, Instituto Nacional de Estadística y Geografía, 2014) and climatic variables that were previously associated with shifting outcomes of plant-rhizosphere interactions and adaptation in *Zea* spp. and other plants (Sawers et al., 2009; Kivlin et al., 2013; O’Brien et al., 2019). Sites ranged 6.6◦C in mean annual temperature (MAT), *>*1100 meters in elevation, and the wettest site received nearly twice the annual precipitation of the driest site (information extracted with raster, Bioclim in R, Hijmans et al., 2005; Hijmans, 2015; R Core Team, 2019, Table S1). In August 2013, we collected 2 kg of teosinte rhizosphere soil from each population (pooled individuals spanning the spatial extent), stored briefly at 4◦C, then sent it for analysis at INIFAP, Laboratorio Nacional de Fertilidad de Suelos y Nutricíon Vegetal. Sites had an *≈*10-fold difference in extractable soil phosphorous (29.7-223 ppm) and potassium (96-1055 ppm), and inorganic nitrogen ranged from 12 to 17.6 ppm. These variables did not shift independently across sites: as MAT increased, so did precipitation, soil water holding capacity, phosphorus, and potassium, but inorganic nitrogen decreased (*ρ* is 0.30, 0.55, 0.41, 0.54, and -0.27, respectively).

In December 2013, after plant senescence and seed set, we collected seeds from 12 different mother plants per population, chosen to span the population spatial extent and have sufficient seed quantity, and stored at 4◦C until use. At the same time, we scored coarse phenology of each population, and collected rhizosphere biota. Approximately 6 liters (4-7 L) of roots and attached soil were collected from plants spanning the whole population at each site. Plants were unearthed and roots lightly shaken, and then roots and loosely-adhering soil were placed in bags, dried at ambient temperature, and stored at 4◦C. To make biota inoculum for each source site, bag contents were homogenized in a blender until root pieces were approximately *≤* 2 cm in length and well mixed with soil. While pooling soil samples within sites can homogenize within site variation, homogenization effects should be unbiased with respect to local adaptation between plants and rhizosphere biota. To characterize abundance of a key rhizosphere microbe in inocula, we extracted arbuscular mycorrhizal spores from homogenized inocula (density gradient method Furlan et al., 1980).

### Testing whether biota from stressful sites provide more general and locally adapted benefits

In May of 2014, we grew seeds from each teosinte population in each of six inoculum treatments: no inoculum, sympatric inoculum (collected from same site, contrasted with “allopatric,” collected from different sites), and inocula from four sites selected from the 10. These treatments ensured that each teosinte population experienced biota from its home site and biota from allopatric sites. The four plant populations from which these selected inocula came received doubled replicates of the sympatric treatment, and three allopatric treatments, while other populations received four allopatric treatments (see Figure S2). Source sites used for the shared biota inocula treatments spanned the range of described environmental variables (Table S1, Figure S2).

We grew sibling seeds from 12 mothers from each of the 10 teosinte populations (120 mothers *×* 6 treatments = 720 plants). We added four drainage holes to 2 L plastic grow bags, and filled with 1.5 L of sterile potting mix (90% sand, 5% perlite 5% vermiculite 0.2% silt, steam sterilized for 4 hours at 90◦C using a PRO-GROW SS60). We inoculated each pot with 50 mL of 4:1 sterilized sand and homogenized inocula (sterilized sand only in uninoculated treatment) just below where seeds were to be placed, and topped with sterilized soil, resulting in a live layer of inocula sandwiched between sterilized soils. As only 0.5% of pot volume is inocula, we expect any non-biotic inocula effects to be minimal relative to biotic effects. We added three seeds from the same maternal plant family to pre-watered pots after scarification with overnight soaking, and thinned to one seedling after germination. Pots were randomly arranged on a bench in a temperature- and humidity-controlled greenhouse in Irapuato, Gto, Mexico (average temperature 23.8◦C during the experiment). We treated plants with Agrimycin and Knack in dual-application one time to prevent caterpillar and spider mite herbivory. We kept pots unfertilized and moist for the first two weeks as most plants germinated, after which we watered and fertilized weekly with 50 mL of Hoagland’s solution adjusted to low phosphorous (100*µ*M). We chose this low nutrient and phosphorous regime to increase stress that rhizosphere interactions could alleviate (Smith et al., 2010), as recommended for tests of COCO (O’Brien et al., 2018).

At 52 days post-germination (dpg), we harvested plants. When many plants were due for harvest on a particular day, we harvested over several days in random order; most plants were harvested within one or two days of 52 dpg (Figure S3; range 29-67 dpg). We measured traits (see below), then quantified a fitness proxy: pre-reproduction vegetative dry biomass, which predicts fitness in the related subspecies *Zea mays* ssp. *parviglumis* (Piperno et al., 2015, as analyzed in O’Brien et al., 2019). We washed plants of adhering soil, split into roots and shoots, dried (*≈* 45◦C until mass stabilized), and weighed.

We related our fitness proxy, biomass, to abiotic environments at plant and biota source sites with linear models (Bayesian methods, MCMCglmm Hadfield, 2010). Our environmental variables included our three soil fertility measures (logged when normality improved), soil water holding capacity, and climatic variables (site mean annual temperature, mean annual precipitation). We explicitly tested whether biota effects on plant fitness are correlated to the environment at their source sites (*E_B_*), whether local adaptation alters these effects (*S*), and whether local adaptation is environment-specific (*E_S_ × S*) by fitting:

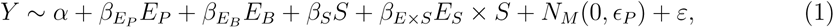

where *β*s are slopes, and *α* is the intercept. Biota source environment effects (*β_E_B__*) may be a combination of species assemblage differences and divergence within rhizosphere species. Sympatric effects (*β_S_* and/or *β_E×S_*) may be plant- or biota-based. If *β_S_* is positive, it may indicate local adaptation of plants to perform better in local biota, or filtering of biota via competitive exclusion or selection that results in biota that better support local plants. If negative, *β_S_* may indicate biota that grow more themselves at the expense of local plants. Significant *β_E×S_* would indicate a strengthening or weakening of local adaptation across abiotic gradients. We define all possible sources of *β_S_* and *β_E×S_* as local adaptation, as all involve a local-genotype dependent effect. As stress decreases with increases in our *E* variables, COCO predicts negative *β_E_B__* (biota from colder, drier, and nutrient-poor sites more beneficial) and negative *β_E×S_* with a positive *β_S_* (biota from colder, drier, and nutrient-poor sites provide even greater benefits to sympatric plants).

We include plant source environment effects (*β_E_P__*), as it is important to account for population effects when testing for local adaptation (Blanquart et al., 2013; O’Brien et al., 2018), which could include genetic differences across populations and transgenerational environment responses (i.e. maternal effects). *E_P_* is a random effect for family (which can shape variation in teosinte, O’Brien et al., 2019, Table S3), and *ε* is error. We fit this model using each environmental variable in turn, removing non-significant terms until DIC (Bayesian verion of AIC, Spiegelhalter et al., 2002) stopped reducing or no non-significant terms remained (terms were removed one at a time, starting with most-complex and least significant based on pMCMC). We report the model for only the best fitting environmental variable, quantifying uncertainty with highest posterior density intervals (HPDI, Bayesian equivalent of confidence intervals, Plummer et al., 2006).

### Testing whether stress gradients and local adaptation shape traits and element profiles

We measured plant traits largely from within the set of previously known adaptive or rhizosphere-influenced traits in teosinte or *Zea mays* subspecies (Kaur et al., 1985; Lauter, 2004; Hufford et al., 2013; López et al., 2011; O’Brien et al., 2019). We recorded germination date, measured height to the highest ligule at five timepoints, and length and width of the second true leaf when expanded. At harvest we measured: final height, stem width (at the first node above the soil), leaf number, and number of stem macrohairs in 1 cm^2^ below the ligule on the edge of the lowest live leaf sheath. Some plants germinated much later than others (14 plants surviving to harvest germinated 30-72 days late). These were excluded from analyses including multivariate trait axes (see below) as they could not be measured for all traits, though they were included for biomass, above. We characterized growth timing by fitting parabolic growth curves to height measurements using days since emergence and the square of days (linear models in R, *height ∼ α* + *β*_1_*days* + *β*_2_*days*^2^). We extracted the coefficient for the squared term (*β*_2_), which separated plants into early (*β*_2_ *<*0 plants grew quickly early, with decreasing growth rate through time, no plants had negative growth), or delayed growers (*β*_2_ *>*0, plants initially grew slowly, and increased growth rate through time, see Figure S3). We sampled the youngest (most apical) fully expanded leaf, dried at 45◦C then processed at Donald Danforth Plant Science Center to quantify plant tissue concentration of 20 elements using inductively coupled plasma mass spectrometry (as in Baxter et al., 2008, ICP-MS, ionomics, see Figure 2 for list).

Most element concentrations and some traits were not normally distributed. We took the natural log when this improved normality, but for phenotypes, we restricted taking the log to only traits where the Shapiro W statistic was *<* 0.9, evaluated in R; after necessary transformations all W were *>* 0.75, Figure S4). For all elements, we included greenhouse and field (see below) samples when evaluating normality. Only arsenic and selenium remained substantially non-normally distributed (best W statistic of Shapiro test *<* 0.75, Figure S5).

We tested for differences between uninoculated plants and plants inoculated with biota. We used linear models (MCMCglmm, in R) for each element or trait. Because there were many elements and traits, we used linear discriminant analysis to explore multivariate differences with inoculation (LDA, package MASS in R, Venables and Ripley, 2002).

We used canonical correlation analysis (with package CCA in R, González et al., 2008) to find the axes of greatest multivariate covariation between traits and elemental profiles, which we interpret as the traits that most likely depend on nutrient provisioning by biota. As our experiment was conducted at low phosphorus, we also explore phosphorus concentrations in particular. Plants may highly mis-express traits under artificial deprivation of soil biota (Partida-Martinez and Heil, 2011; Hubbard et al., 2019; O’Brien, 2019), so we restricted this analysis to inoculated plants, though we projected uninoculated plants onto resulting CCA axes for comparison. We evaluated links between composite axes and fitness using linear models (*fitness ∼ axis*), fit with MCMCglmm in R. To compare to results for local adaptation, we performed the same linear model analysis as for biomass (above) on the first two CCA axes and the traits most strongly correlated to them (strongest loadings), as well as leaf tissue phosphorus, due to expected links to a key rhizosphere component (AMF, Smith et al., 2010). To contrast these results with multivariate axes of traits and elemental profiles that may not be linked to each other (or to fitness, see Figure S12), we further extended the analysis to the previously described LDA axes, and the first axes of separate principal components analysis for traits and elemental profiles (see Table S4).

### Testing whether biota provide greater benefits at stressful sites and retain effects on traits and elements

Because predictions of COCO rest on increasing benefits of biota at stressful sites, and some adaptive benefits may be conditional on local environments, we evaluated relationships between environment, elemental profiles, size traits, and rhizosphere colonization at field sites. We focused on one important rhizosphere component: arbuscular mycorrhizal fungi (AMF, Smith and Read, 2008).

During August 2013 collections, we quantified differences in field teosinte plants across the sites. We measured 20 plants per population (spanning the spatial extent) for height to the highest ligule, and stem width at the first node visible above the soil (only these traits could be measured in the field). We sampled the penultimate leaf (to avoid leaves still expanding), stored in paper envelopes, and included these in ICP-MS analyses described above. We also took a sample of mixed roots from throughout the upper 15 cm of the root system, transported from the field in 8 mL tubes of 10 % KOH, which we scored for AMF arbuscules using standard methods (McGonigle et al., 1990), modified with less toxic alternatives. Briefly, we left roots in their field KOH solutions to clear (5-10 days), placed subsamples in histology cassettes, rinsed with deionized water, acidified in acetic acid (5%) for 2-3 hours, boiled in 5% acetic acid and 5% pen-ink (Parker, Quink Black-Blue Waterproof) for 3-5 minutes (until roots take up the ink), and rinsed once with deoinized water. We mounted stained roots on slides in corn syrup (Karo brand), and scored approximately 60 intersections for arbuscules with brightfield illumination microscopy (Vierheilig et al., 2005).

We projected field elemental profiles onto significant CCA axes we calculated from greenhouse data above. We selected the best performing environmental variable (present in the most best models, *β_E_*) from the greenhouse data to use in the field models, and we tested if projected field elemental profiles, field tissue phosphorus, or field-measured traits, suggested that effects of AMF (*β_C_*) increase at more stressful sites (*β_E×C_*):

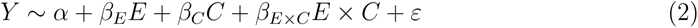

We fit the full model first and removed non-significant terms one at a time (as above, analogous analysis for LDA and PCA in Table S5).

## Results

### Only the COCO prediction of increased local benefits for plants and biota from stressful sites is supported

For biomass (our fitness proxy), we found that the best fitting plant and biota source variable was mean annual temperature (MAT, see also Figure S6). COCO predicted that biota from stressful cold sites should be the most generally mutualistic. Contrary to these predictions, overall, association with biota from warmer, less stressful sites increased plant biomass in the greenhouse (Figure 1, *β_E_B__*, Table 1). However, sympatric combinations of plants and biota produced greater plant biomass than expected from the effects of plant source and biota source on biomass (*β_S_*, Table 1), suggesting benefits from local adaptation. In line with COCO predictions for local mutualistic adaptation, benefits of local adaptation were stronger for plants from colder sites: the interaction effect between MAT and sympatry (*β_E×S_*) eroded the effect of sympatry (Table 1), reflecting that teosinte from colder sites paired with sympatric biota more strongly exceeded expectations for biomass when excluding sympatric terms (Figure 1, right).

**Figure 1:**
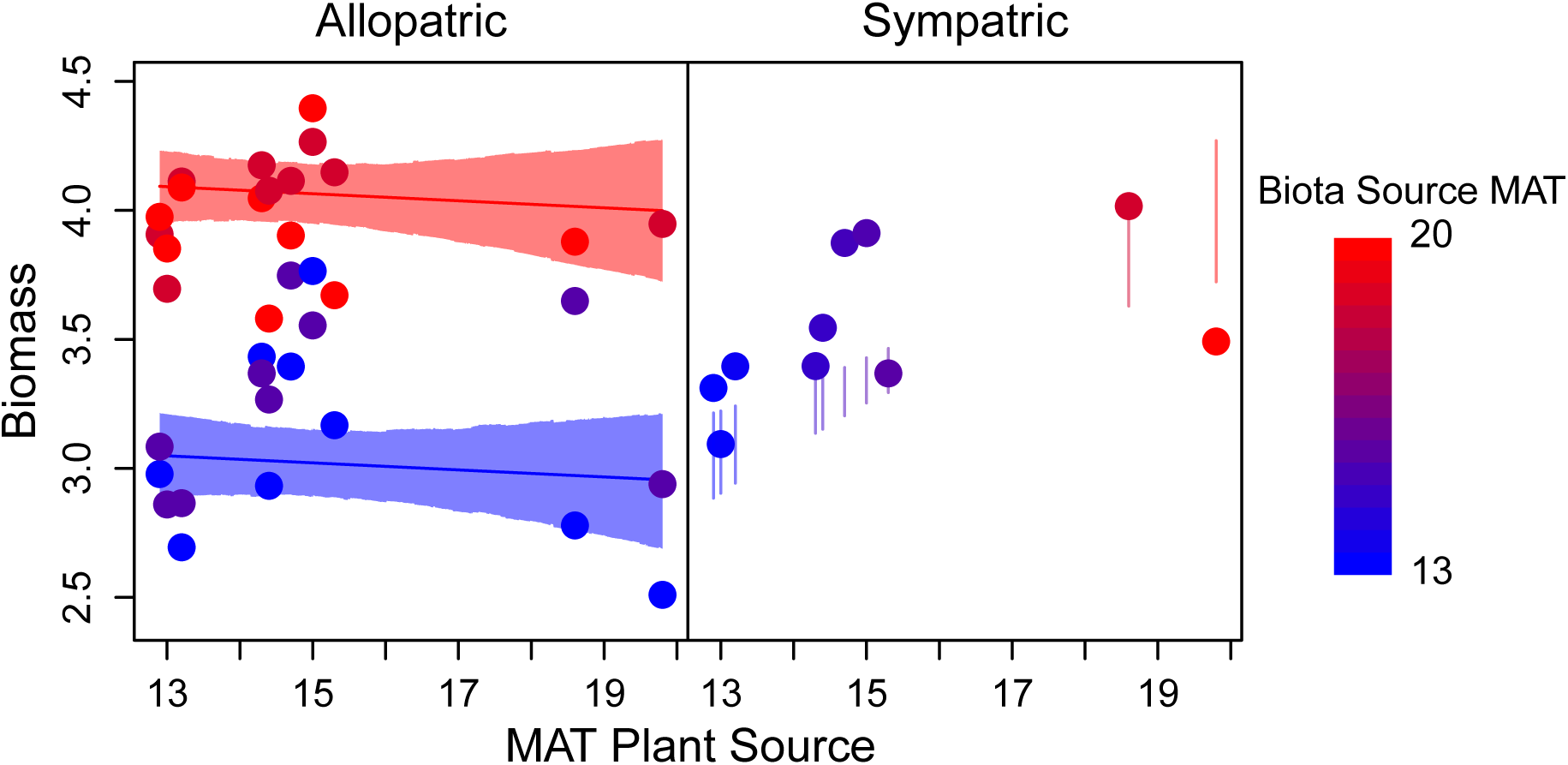
Mean biomass for each combination of plant and rhizosphere biota sources (points) plotted against the mean annual temperature (MAT *^◦^*C) of the plant source site. Left panel: allopatric combinations show greater generalized mutualistic benefits from biota from warmer sites; non-overlapping model expectations (lines, 95% HPDI for the mean in shaded intervals) between plants grown with biota from the warmest (red) or coldest (blue) sites (effect of plant source MAT n.s.). Right panel: sympatric combinations (means, points) of teosinte and biota from colder sites show greater local, sympatric benefits (fall above vertical lines: 95% HPDI model expectations for these points excluding sympatric effects, *β_S_* and *β_E×S_*). Point color indicates MAT at the source site of the inoculated biota for both panels (redder = warmer).

**Figure 2:**
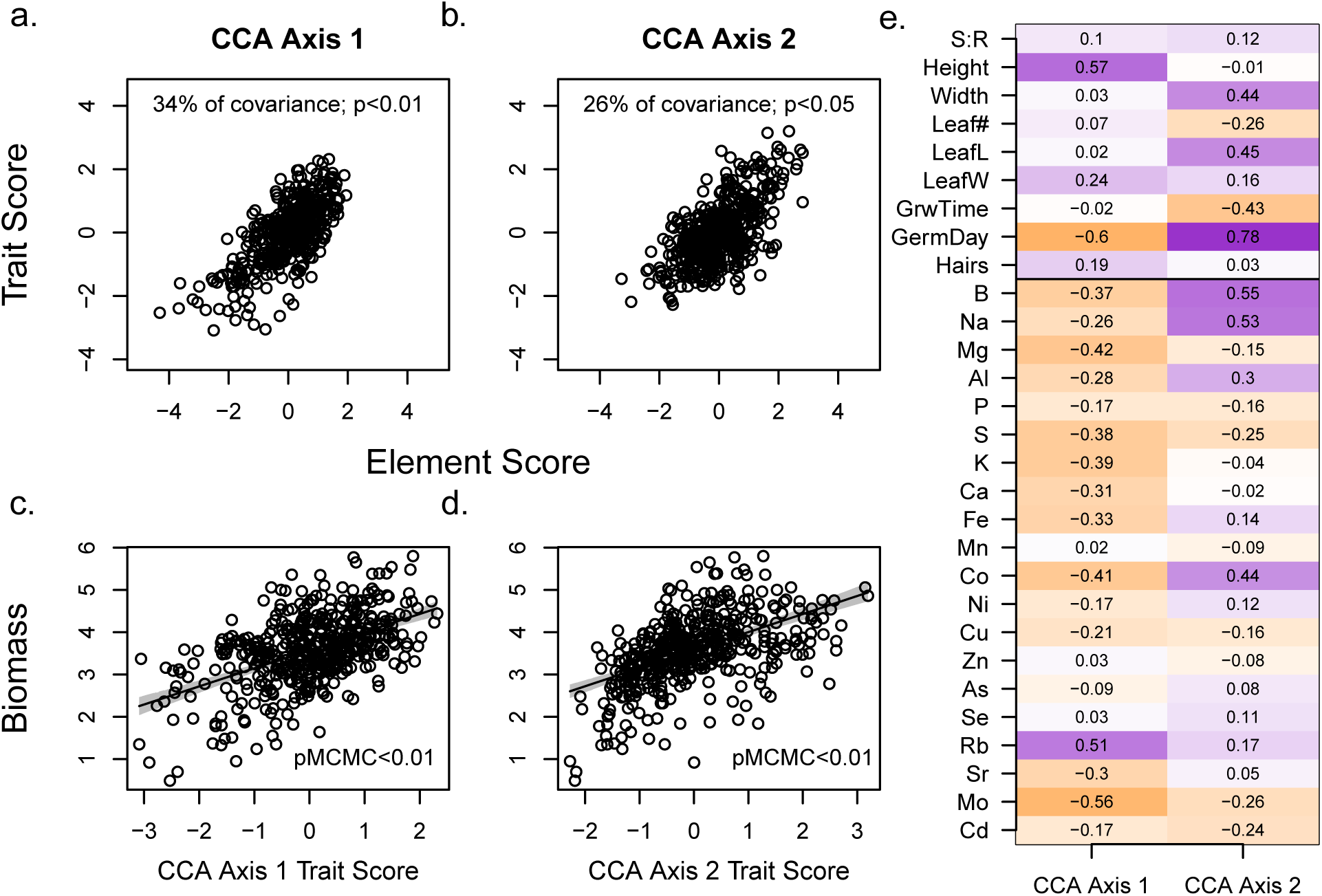
Two significant (as in Tabachnick et al., 2007) multivariate axes of covariation between traits and elements in greenhouse plants were identified by CCA, depicted by plotting trait and element scores for each axis (a and b, axis 1 and 2, respectively). Both multivariate axes were strongly positively correlated to biomass, for trait (c and d, for axis 1 and 2, re-spectively) and element scores (not shown, both pMCMC*<*0.01). (e) shows full multivariate correlations (equivalently, “loadings”) of trait (top) and elements (bottom) on CCA axes. Stronger positive correlations are in darker purple, stronger negative correlations in darker orange. Standard abbreviations for elements, GermDay = natural log of day of germination, lnHair = natural log of stem hairs per cm^2^, S:R = shoot:root ratio, Width = stem width, LeafL = leaf length, LeafW = leaf width, GrwTime = higher values indicate a delay in peak growth rate. See Figure S9.

**Table 1:** Biota and plant source effects estimated by best models for plant fitness (biomass), plant tissue phosphorus, and trait scores on first and second CCA axes.^1^

|  | *parameter* | biomass | ln(phosphorus) | CC1 Trait | CC2 Trait |
| --- | --- | --- | --- | --- | --- |
| Intercept | $a$ | 1.27** | 7.13** | 0.92. | -3.97** |
| Biota source Env. | $\beta_{EB}$ | 0.15** | – | 0.16** | 0.11** |
| Plant source Env. | $\beta_{EP}$ | -0.014 | -0.017* | -0.24** | 0.15** |
| Sympatry | $\beta_S$ | 1.32* | -0.44* | 1.68** | 1.53* |
| Sympatry × Env. | $\beta_{E \times S}$ | -0.077* | 0.025* | -0.10** | -0.096* |
| Env. in best model |  | MAT | MAT | MAT | MAT |

### Inoculation increases phosphorus, biomass and affects elemental profiles, traits

Inoculation with biota had the expected effects of increasing biomass and tissue phosphorus relative to levels in uninoculated siblings. Inoculated plants were over 30% larger (average 3.50 and 2.59 grams total biomass, *±* 0.04 and 0.09 SE, pMCMC *<* 0.05, see Table S3). Tissue concentrations of phosphorus were nearly double in inoculated plants (977 µg g*^−^*^1^ versus 569 µg g*^−^*^1^, standard error 11.8 and 12.1, respectively), but were still below levels for ideal plant growth (3000 µg g*^−^*^1^ Marschner, 2011, Figure S13). Beyond phosphorus, some greenhouse plants (and field plants) had concentrations in their tissues potentially signalling deficiency (magnesium and molybdenum) or toxicity (sodium, Figure S13, Marschner, 2011).

Our multivariate LDA distinguished the elemental profiles of inoculated plants from those in the sterilized treatment (successful assignment 95% overall) primarily based on tissue phosphorous (Figure S7). LDA of trait data poorly distinguished plants growing with live biota from uninoculated plants (predicted only 5% of uninoculated plants) but plants in live biota had wider stems (6.2 and 5.5 mm, *±* 0.07 and 0.13 SE) and grew earlier (Figure S8, and Table S3, both differences pMCMC *<* 0.05). Most plant traits and element concentrations had significant correlations across inoculated and uninoculated siblings, indicating contributions from maternal environments, non-plastic genetic differences among families or populations, or similar (Table S3).

### Co-varying axes of elemental profiles and traits link to fitness

Because we expected plant nutrition and plant traits to be causally linked, we employed canonical correlation analysis (CCA) to identify the strongest axes of covariation between them. The first two axes explained significant covariation between traits and elements (34% and 26%, chi-squared *p <* 0.01 and 0.05, respectively, Figure 2a-b,e), and explained moderate portions of variance within traits (9% and 15%) and tissue elements (10% and 6%, respectively). For ease of interpretation, we flipped loading signs on the first CCA axis; this does not change results.

CCA axes identify highly multivariate relationships that may not be easily simplified into components, however, we identified several patterns. Briefly, on the first CCA axis, for a given concentration of rubidium, plants that had decreased tissue concentrations of molybdenum, cobalt, magnesium, potassium, and the majority of other elements were taller and also germinated earlier (Figure 2e, see Figure S9 for partial axis visualization). This axis may relate to potassium nutrition, as it is orthogonal to the well-known positive correlation between potassium and rubidium (Läuchli and Epstein, 1970, Figure S9). On the second axis, plants with elevated boron, sodium, and cobalt were linked to plants that germinated later but had wider stems, longer leaves and earlier timing of maximum growth rate (negative values for growth timing, Figure 2, see Figure S9 for partial axis visualization). Projected scores for uninoculated plants were lower than inoculated plants on the first and second axes (Figure S10, i.e. because they were smaller and had higher tissue concentrations of most elements excepting phosphorus, Figure S7, Table S3).

We expected that traits and elemental profiles would link to fitness. In the greenhouse, biomass was strongly correlated to CCA axes of elements and traits, as would be expected if relationships were causal or had underlying shared causes (Figures 2c-d,S12, *ρ >*0.5 for CCA axis 1). Instead, for multivariate analyses agnostic to trait links, trait and element axes are not correlated to each other (Figure S11). Correlations to biomass were generally weaker or even anti-predictive for these axes and phosphorus (Figure S12). For example, while both phosphorus and biomass increased with inoculation, phosphorus was negatively correlated to biomass among inoculated plants (*ρ* -0.21).

### Local adaptation and source of plants, biota co-affect traits and elemental profiles

Using linear models, we tested whether plant tissue phosphorus and linked axes of traits and elemental profiles differed across the environment of plant and biota sources. As seen for biomass, mean annual temperature (MAT) was the best fitting plant and biota source variable for these response variables (Table 1, see Table S4 for element score best models).

We found mixed effects of locally matched plants and biota on tissue phosphorus and co-varying traits and elements. Plants from colder sites had more tissue phosphorus, but the only difference across biota was that teosinte from colder sites growing with sympatric biota had relatively lower tissue phosphorus (Figure 3a, Table 1), both trends likely reflecting elevated tissue phosphorus of smaller inoculated plants (see above). Plants sourced from colder sites had increased values on the first CCA axis for both traits and elemental profiles (Figure 3b, taller plants that also germinate early, and have high tissue rubidium relative to molybdenum, potassium, and most other elements), but plants growing in biota sourced from colder sites had decreased values for both traits and elemental profiles on the first CCA axis (shorter plants that also had later germination and opposite elemental profile shifts, see also Figure S9). Sympatric biota moved scores on the axis in the same direction as biota from warmer environments (positive sign of *β_E_B__* matches *β_S_* Tables 1, S4), and, as seen for biomass, the strength of the sympatric effect decayed for plants and biota from warmer sites (negative *β_E×S_*, Tables 1, S4).

**Figure 3:**
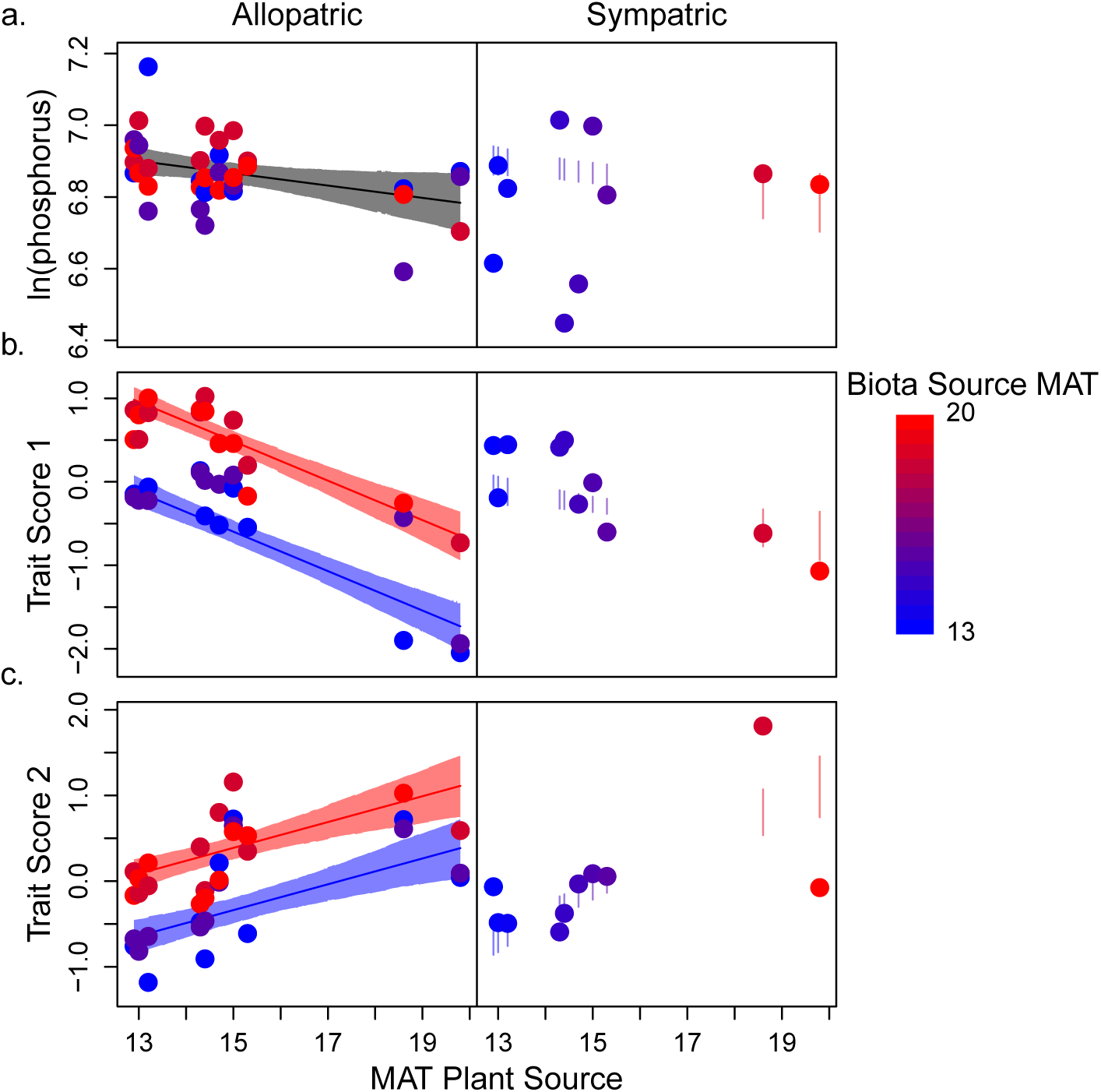
Mean tissue phosphorus (a, natural log) and CCA axis trait scores (b, c) for each combination of plant and biota sources (points) plotted against the mean annual temperature (MAT *^◦^*C) of the plant source site. Point color indicates MAT at rhizosphere biota source site (redder = warmer). Left panels: observations and model expectations for allopatric treatments (lines, mean; shaded region 95% HPDI), separating predictions for plants grown with biota from the warmest (red) or coldest (blue) sites (except for phosphorus; *β_E_B__* is n.s). Right panels: observations for sympatric combinations. Vertical lines give 95% HPDI for model expectations for means omitting sympatric effects, as in Figure 1. Observed means outside this interval suggest local adaptation. See Table 1.

While plant source MAT and biota source MAT had opposite effects on the first CCA axis, they had aligned effects on the second (*β_E_B__* & *β_E_P__ >* 0, Figure 3c). This signals that plants from warmer sites and plants growing with biota from warmer sites had later germination combined with earlier growth, and wider (but not taller) stems, as well as relatively higher concentrations of elements that load positively on CCA axis 2 (boron, sodium, and cobalt, see Figure S9). Sympatric biota shifted trait scores in the same direction as plants and biota from warmer environments, but sympatric effects decayed for plants from warmer environments (*β_S_ >* 0, *β_E×S_ <* 0, Table 1, Figure 3c, element score best model differs, Table S4). Similar results between the best models for CCA axes and best model for biomass reflect positive correlations between both CCA axes and biomass in the greenhouse (Figure 2, Figure S12).

### Field data suggest increased benefits of biota in cold sites

COCO predictions rest on increased benefits of biota to plants at stressful sites. Both height and stem width increased with colonization at colder sites, but weakly decreased with colonization at warmer sites (interaction pMCMC *<* 0.01, Table 2, Figure 4b,d). This is consistent with greater benefits of biota at colder sites, and with positive effects of sympatric biota for plants from colder field sites (in the field, all plants associate with sympatric biota). In field plants, the projections onto the first CCA axis are negatively correlated to mean annual temperature, concordant with plant source effects observed in the greenhouse (less concentrated elements, taller plants at colder sites, slope pMCMC *<* 0.01, Tables 1, 2, S5, Figures 3b, 4a,c), and therefore opposite to biota source effects observed in the greenhouse. However, field plants from colder sites are only taller when colonized by mycorrhizal fungi (Figure 4b, Table 2). Field plant scores on both this axis and the LDA for elemental profiles may reflect greater biotic inoculation: field plants are shifted away from uninoculated plant scores in the greenhouse, in the same direction as, and exceeding inoculated greenhouse plants (Figures S7, S10; Table 2). Compared to greenhouse plants, field plants had lower tissue concentrations of more than half of the elements, but higher concentrations of rubidium, Figure S13), and were indeed taller: average height was 80*±*4.5 and 35*±*0.4 cm in the field and greenhouse respectively (means*±*SE, pMCMC *<* 0.01).

**Figure 4:**
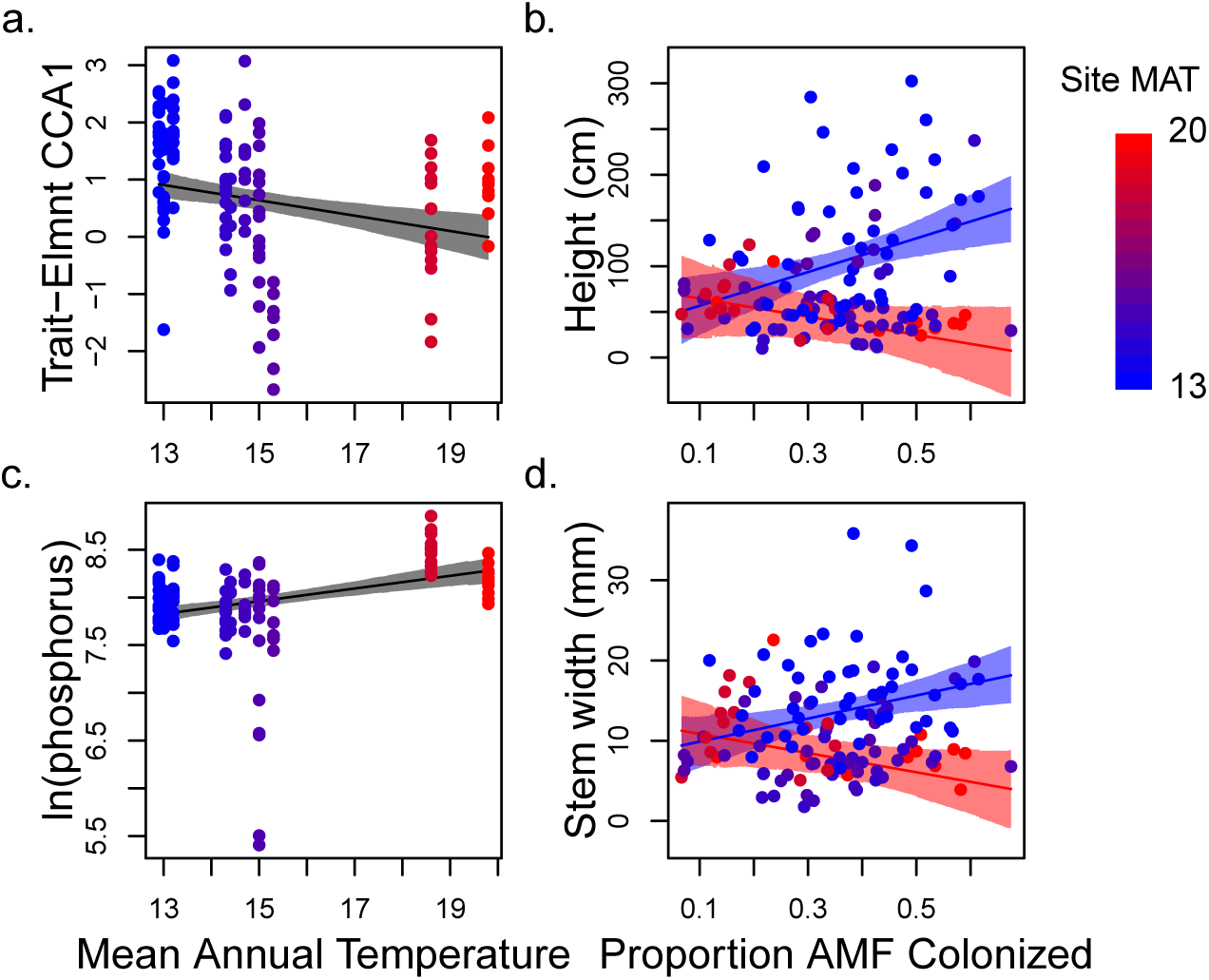
Field plant projected element scores on CCA axis 1 (a), tissue phosphorus (c, natural log of *µ*g g*^−^*^1^ dry weight), and measured traits (b, d), plotted against either site mean annual temperature (MAT◦C, a & c) or proportion of field roots colonized by AMF (b, d) when AMF effects were in best models. Color indicates MAT of the site (redder = warmer). Predictions (lines) and 95% HPDI intervals (shading) are shown for the mean across MAT of the sites in gray (a, c), or for the mean across the proportion of root length colonized by AMF, split into expectations for the warmest and coldest sites (red and blue, respectively) for height (b), and stem width (d).

**Table 2:** Best models for the projected element scores of the field plants onto the CCA axes calculated from greenhouse plants.^2^

| | Intercept | $\beta_{MAT}$ | $\beta_C$ | $\beta_{MAT \times C}$ |
| --- | --- | --- | --- | --- |
| **CC1 Element-score** | 2.65** | -0.13** | — | — |
| **ln(phosphorus)** | -6.96** | 0.067** | — | — |
| **Height** | -27.0 | 5.10 | 710.6** | -40.9** |
| **CC2 Element-score** | -4.28* | -0.33** | -8.87. | 0.70* |
| **Stem width** | 1.78 | 0.052 | 63.4** | -0.38* |

On the second CCA axis (associated with phenology and weakly with wider stems), projections for field plants again depended on AMF colonization. More colonized plants were higher on this axis, in the direction of the main sympatric biota greenhouse effect, but patterns across sites were inconsistent with greenhouse patterns. While we were not able to measure phenology in the field, stem width patterns partially match changes in the axis: like greenhouse plants from colder sites, field plants from colder sites indeed had wider stems, especially with AMF colonization (Figure 4d, Table 2), but field plants have much wider stems than inoculated greenhouse plants (11*±*0.43 and 6.1*±*0.06 mm, respectively, pMCMC *<* 0.01) despite equivalent scores on this axis (pMCMC *>* 0.05, Figure S10).

While the CCA axes showed strongly or marginally similar trends in the field, trends for PCA projections (agnostic to trait-element links) did not show any pattern across field sites, despite variation in greenhouse plant PCA scores across source sites (Tables S5, S4), suggesting that the linked trait-element CCA axes are better, though imperfect, predictors of field patterns.

## Discussion

Abiotic environments play a consistent role in structuring local adaptation in plants and other species, but local adapation to biotic environments may be highly variable (Briscoe Runquist et al., 2020; Hargreaves et al., 2020). Rampant shifts in the outcomes of species interactions across environmental conditions (Bronstein, 1994; Bertness and Callaway, 1994; Chamber-lain et al., 2014; He and Bertness, 2014) perhaps underlie this variable strength of local adaptation (e.g. Thompson, 2005; Bronstein, 2009; O’Brien et al., 2018). Like most species interactions, plant interactions with rhizosphere biota vary substantially in both outcomes (Berg and Smalla, 2009; Smith and Read, 2008; Anacker et al., 2014) and degree of local adaptation (Rúa et al., 2016). Further, plant traits are influenced by rhizosphere microbes that they associate with (Friesen et al., 2011), especially through changes to plant nutrition (Desbrosses and Stougaard, 2011; Paszkowski and Gutjahr, 2013; Lu et al., 2018), suggesting that adaptive changes in plant-microbiome interactions may alter the concentrations of elements in plant tissues. We investigated the influence of rhizosphere biota on plant fitness, trait expression, and elemental profiles in teosinte within the context of correlated environmental gradients of both climate and soil fertility. We found that changes in fitness (via a proxy of biomass) and traits were linked to changes in elemental profiles and were affected by the source of the rhizosphere biota and the plant population it was paired with.

### Local adaptation is strengthened in cold sites

Plants from colder sites derived greater specific benefits from their local biota, matching one of our predictions based on COCO (O’Brien et al., 2018): increased local adaptation between plants and biota from colder sites, which we presume are more stressful (Figure 1, using biomass as a proxy for fitness, see Methods). However, biota from colder sites produced less fit plants, in contrast to our other prediction: that biota from stressful sites would provide greater generalized benefits across plants. This prediction of COCO relies on at least some benefits provided by biota to plants being independent of plant genotype and environment, i.e. if some microbes always provide more phosphorus than others, this would likely be beneficial across all hosts and environments. However, our results are consistent with most benefits of rhizosphere biota being either host-plant- or environment-dependent: i.e. there may be little variation in benefits from rhizosphere biota that is not context dependent, and thus limited potential for the evolution of generalized benefits.

Recent experimental work has suggested that more benefits from plant-microbe interactions may derive from local adaptation and genotype-dependent effects than previously thought (Batstone et al., 2020; Ramírez-Flores et al., 2020). Other studies have also suggested that benefits provided by biota to plants are greatest when experimental conditions match the environment to which the biota are adapted (Johnson et al., 2010; Lau and Lennon, 2012), and mean greenhouse temperature during our experiment was closer to mean annual temperature of our warmest sites (Table S1, Methods). This prevalence of host- and environment-dependent effects suggests that efforts to leverage and manipulate organisms in the plant rhizosphere for increased resilience to abiotic stress in agricultural crops (e.g. Bouwmeester et al., 2019), must tailor solutions to specific sites and cultivars.

Indeed, the existence of environment-dependent benefits is an assumption of COCO to begin with. Given that we expect benefits of rhizosphere biota to increase in cold, drought, and infertile conditions, the benefits we observed in the greenhouse may have been underestimates, especially at cold sites. We used field-measured traits and links between traits, elements, and fitness in the greenhouse as a window into benefits of rhizosphere biota in the field. In the field, multivariate patterns across sites for elemental profiles and traits matched only patterns across plant source sites and sympatric effects observed in the greenhouse. Both field and greenhouse plants from colder sites, especially those paired with sympatric biota (greenhouse) or more colonized by AMF (field), were taller and higher on CCA axis 1 (Figures 3b, 4a). We observed opposite relationships between teosinte size and mycorrhizal colonization between warmer (slope weakly negative) and colder (slope strongly positive) field sites (Figure 4b,d, stem width and height), consistent with increased benefits of biota at cold sites, with previous work showing that maize benefits from mycorrhizae increase in the cold (Zhu et al., 2009), and with increased benefits of these biota in sympatric contexts as observed in the greenhouse. Patterns in the field are thus consistent with greater benefits of biota from and at cold sites.

### Plants and biota have both opposing and aligned effects on traits across environmental gradients

Mechanistically, local adaptation to abiotic environments or biotic interactions must ultimately be based on genetic differences in the expression of traits, yet biotic interactions themselves can alter the expression of traits. For example, microbiomes shape traits from obesity to life history in their animal hosts (Turnbaugh et al., 2008; Gould et al., 2018), and plant-microbiomes shape a comprehensive range of vegetative and floral traits in plants (Friesen et al., 2011; Rebolleda-Gómez et al., 2019). If one species’ influence on another species’ phenotype feeds back to affect its own fitness, selection will shape any genetic variation in the first species affecting traits in the second (so-called ‘extended’ phenotypes Dawkins et al., 1982; Rebolleda-Gómez et al., 2019; O’Brien et al., 2021). Indeed, reciprocal feedbacks of traits on the fitness of interacting species is a condition of co-evolution and likely to be common (Thompson, 2005). Perhaps unsurprisingly, a growing number of examples document evolving extended phenotypes (Lau and Lennon, 2012; Panke-Buisse et al., 2015; Rudman et al., 2019) with the implication that extended phenotypes could contribute substantially to local adaptation or local co-adaptation between interacting species.

Here, we observed that the effects of biota and plant source on plant fitness, elemental profiles, and phenotypes sometimes opposed, and sometimes matched each other. Plants from cold sites were relatively taller with earlier germination, and tissue concentrations of most elements that were low relative to rubidium levels, but biota from cold sites produced relatively shorter and later-germinating plants that had opposite shifts in elemental profiles (Figures 2,3b, Figure S9). However, the second major axis of trait co-variation instead shows similar, or reinforcing effects of plants and rhizosphere biota from the same site. Opposing or correlated impacts on trait values between hosts and microbes have been theoretically linked to the evolution of extended phenotypes under conflicting or identical trait optima, respectively (O’Brien et al., 2021), suggesting that plants and microbes may have different fitness optima across sites for traits on the first CCA axis, but similar optima for traits on the second axis.

### Potential mechanisms of links between elemental profiles and traits

We observed that variation in fitness (biomass) was shaped by the influence of biota and climate on linked, multivariate axes of plant phenotypes and elemental profiles (CCA, Figures 1-3). These linked axes primarily included connections between elements and size (height, stem width) or elements and phenology (germination, timing of peak growth).

Phenology is an important trait for ecological adaptation in both maize and teosinte, with colder high elevation populations flowering earlier and having less seed dormancy (Rodríguez et al., 2006; López et al., 2011; Navarro et al., 2017). Elemental profiles can signal physiological variation (Baxter et al., 2008), but we cannot ascribe causality to particular elements here. However, several elements and traits that load heavily onto the CCA axes match with known links between phenology and nutrition, and may be worth further investigation. Eelays in germination in teosinte from, or growing with biota from, warmer sites were associated with multivariate shifts in elements loading strongly onto CCA axes 1 (higher rubidium relative to the concentrations molybdenum and most other elements) and 2 (including simultaneously increased sodium and boron, Figures 2, 3b-c,S9). Only sodium was at levels expected to be limiting (Figure S13, Maron et al., 2014). Sodium toxicity has been previously linked to inhibited germination in maize, and maize tolerance to sodium can be impacted by rhizosphere biota (Farooq et al., 2015). Likewise, multivariate increases in elements loading positively onto CCA axis 2 (primarily boron and sodium) was associated with plants from, or growing with biota from, warmer sites that had an earlier burst of growth (Figures 1, 3c, S9). Boron is required in large amounts by reproductive tissues of maize (Lordkaew et al., 2011; Marschner, 2011), and precocious flowering can be favored by sodium stress (Farooq et al., 2015), suggesting possible links between sodium, boron, growth timing and flowering in teosinte. In the field, teosinte plants in warmer sites complete flowering earlier (Table S2), and a boron transporter was implicated in adaptation to different climatic environments in teosinte (Pyhäjärvi et al., 2013).

We expected mycorrhizal fungi to drive some linked changes in elemental profiles and size traits, as phosphorus deficiency in our experiment should have enhanced both their colonization of teosinte roots and benefits from phosphorus provided (Smith et al., 2010). Indeed phosphorus and rubidium (which can also indicate increased AMF colonization, Hawkes and Casper, 2002) increased significantly from uninoculated to inoculated plants (Table S3). However, phosphorus patterns in the field (Figure 4c) do not match with spore counts or colonization rates (Table S2, Figures 4b,d, S13), and other changes in our leaf elemental profiles only partially reflect changes observed in maize profiles following inoculation with mycorrhizal fungi (Kothari et al., 1990; Ramírez-Flores et al., 2017). Trade-offs that we observed between small plants with more concentrated elements versus plants that grow larger despite low, or even deficient concentrations of elements (CCA axis 1, Figures 1, S9, S13) could instead be driven by other microbes: a wide array of root-associated bacteria can synthesize, metabolize or interfere with plant hormones (Duca et al., 2014; Gamalero and Glick, 2015) and many soil bacteria alter plant-available nitrogen or phosphorus (Bulgarelli et al., 2013). Indeed, the rhizosphere biota we manipulated here certainly include microbes beyond mycorrhizal fungi.

## Conclusions

Our results highlight the co-influence of abiotic and biotic factors on plant phenotypes. We observed that environment patterned the extent of local adaptation between plants and rhizosphere biota, and the effects of plant-biota interactions on phenotypes. Going a step further, we also know that rhizosphere community composition and function commonly shift across climatic gradients (Veen et al., 2017; Van Nuland et al., 2017; Praeg et al., 2019; Karray et al., 2020; Vieira et al., 2020). As species colonize new habitats in response to global change, the turnover from locally adapted to novel species interactions may drive unexpected phenotypic changes and have implications for successful range shifts.

## Acknowledgements

We would like to thank members of the Eguiarte laboratory for help with collections in the field, Maŕıa Rocio Martinez Villalpando, Carlos Fabián de la Cruz, Abenamar Gordillo Hidalgo, Dario Alavez & Arturo Chávez for greenhouse help. The project was supported by UC MEXUS, the UC Davis Center for Population Biology, GSR fellowship from the UC Davis Department of Plant Sciences, and NSF GRFP DGE-1148897 funding to AMO; USDA Hatch project CA-D-PLS-2066-H and NSF grant IOS-0922703 to JRI; and NSF grant DEB-0919559 to SYS. LEE and JGP work was funded by grants CB2011/167826 (CONACYT Investigación Científica Básica), M12/A03 ECOS Nord France (CONACYT-ANUIES 207571), and by institutional funding from the Institute of Ecology, Universidad Nacional Autónoma de México. Ionomic profiling and IB were supported by the United States Department of Agriculture-Agricultural Research Service. We would also like to thank Graham Coop and Joanna Schmitt for helpful discussions during experiment planning.

## Authorship Contributions

All authors contributed substantially to the design of the study, provisioning of materials, and revising of the manuscript. AMO proposed the study together with JRI and SYS. AMO collected the data, performed analyses and provided the first draft of the manuscript. LEE logistically supported and advised the fieldwork, which was conducted by AMO and JGP. RJHS logistically supported and advised the greenhouse work, which was conducted by AMO. IB advised on sampling design for ionomics, and extracted ionomics data.

## Data availability

All scripts and datafiles are on Github (https://github.com/amob/AO-1) and will be made public at the time of publication; data will additionally be made available on figshare.

---

**Figure S1:**
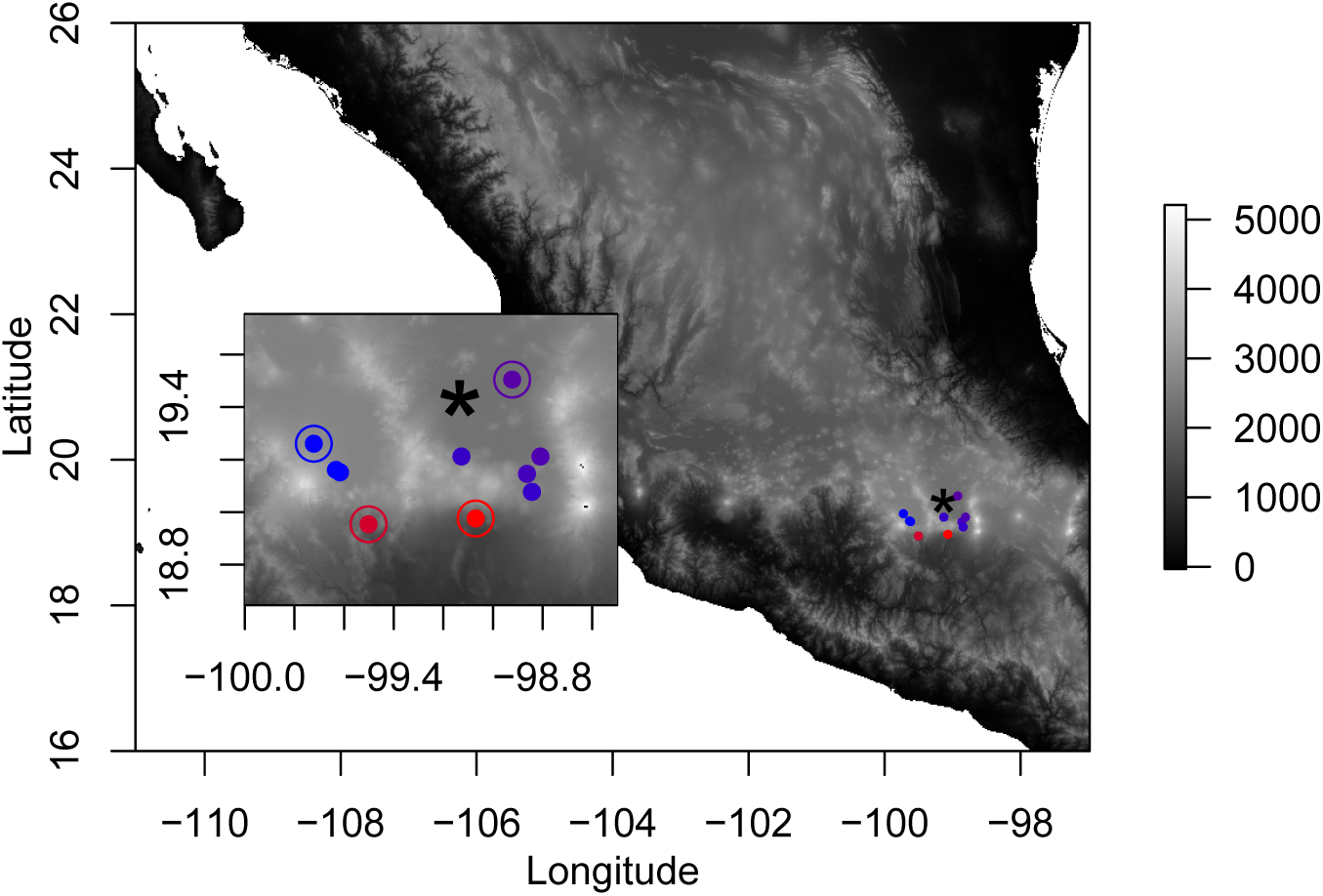
Locations of source sites for teosinte and rhizosphere biota (points), with respect to Mexico City (asterisk), and elevation (color scale, meters above sea level). Inset shows zoom for detail of geographic features around sites, as well as circling the sites from which rhizosphere biota was applied to all plants (see Figure S2, and O’Brien et al., 2019; elevation data from Hijmans et al., 2005).

**Figure S2:**
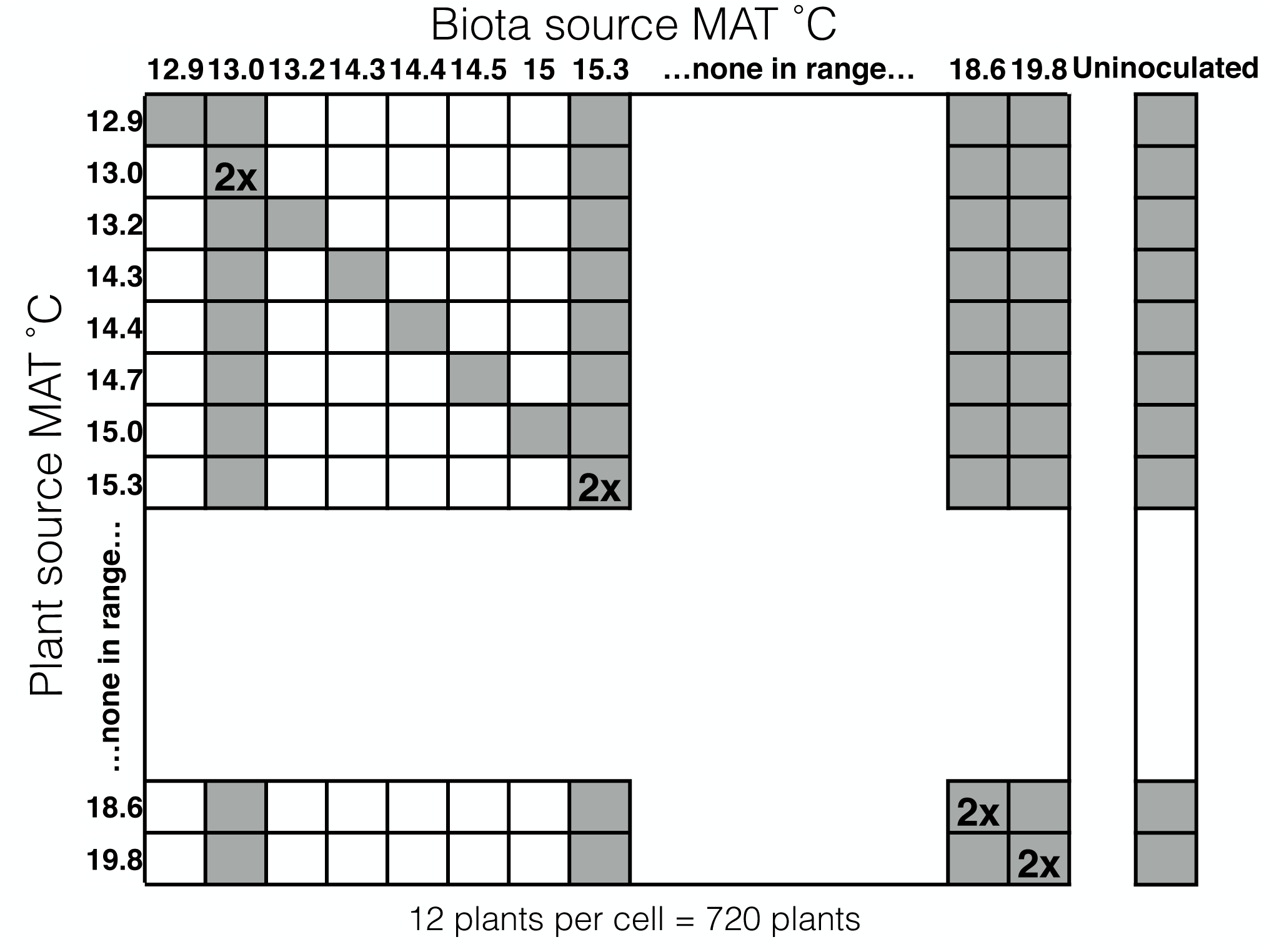
Schematic of experimental design. Outlined columns represent biota sources, outlined rows represent teosinte seed sources. MAT of the site of collection is given for each source. Blank areas represent a significant gap in MAT of sampled sources. Treatments included in the experiment are filled in grey squares and represent 12 pots (one pot per each maternal plant in the field from which seeds were collected), and “2x” denotes double the number of experimental pots (2 pots per field maternal plant, or 24 pots total). All plant populations were also grown uninoculated.

**Figure S3:**
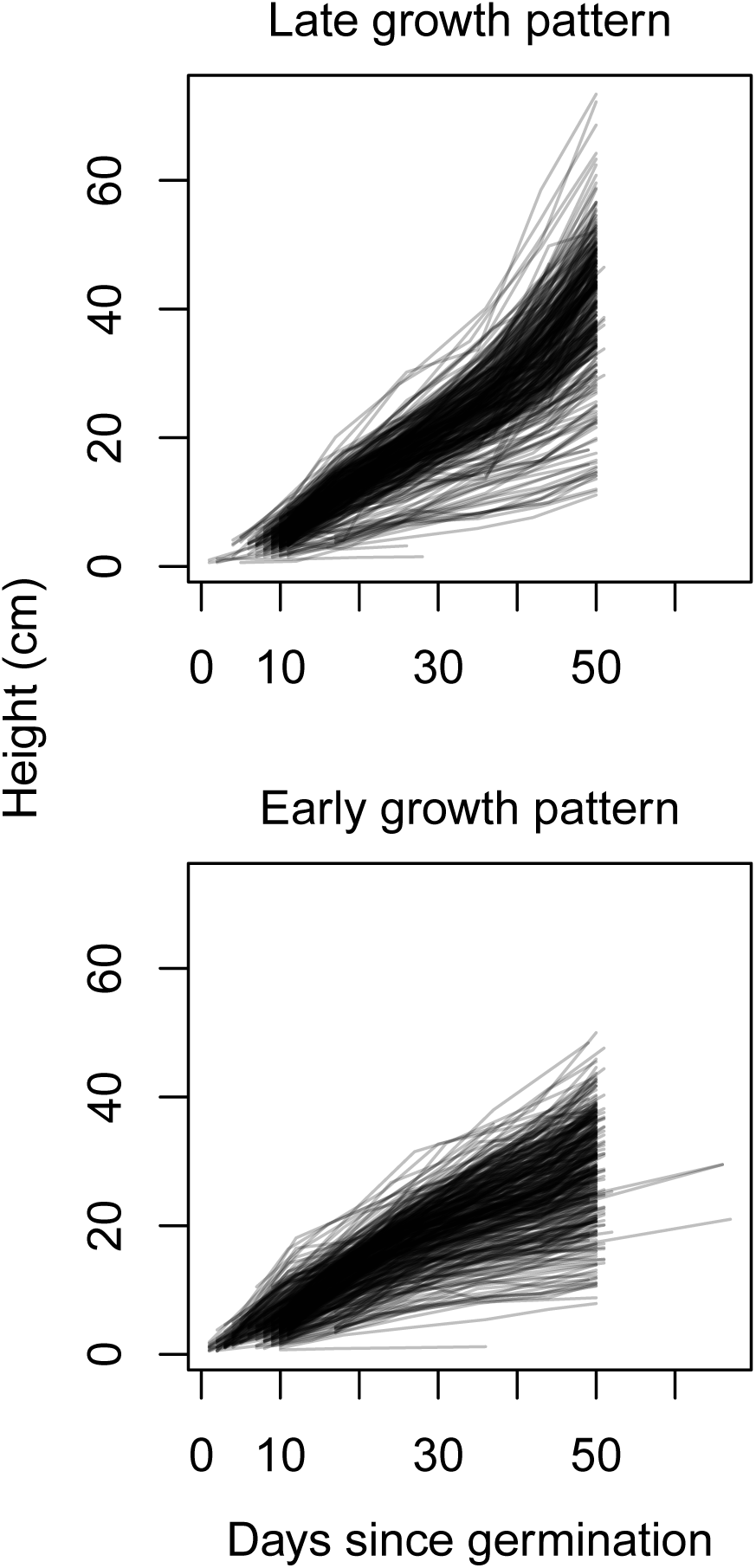
Time series of height through time for all measured plants. Plants fall into either delayed growth pattern (posiive squared term in fitted parabola, top) or early growth pattern (negative squared term in fitted parabola, bottom), see Methods text.

**Figure S4:**
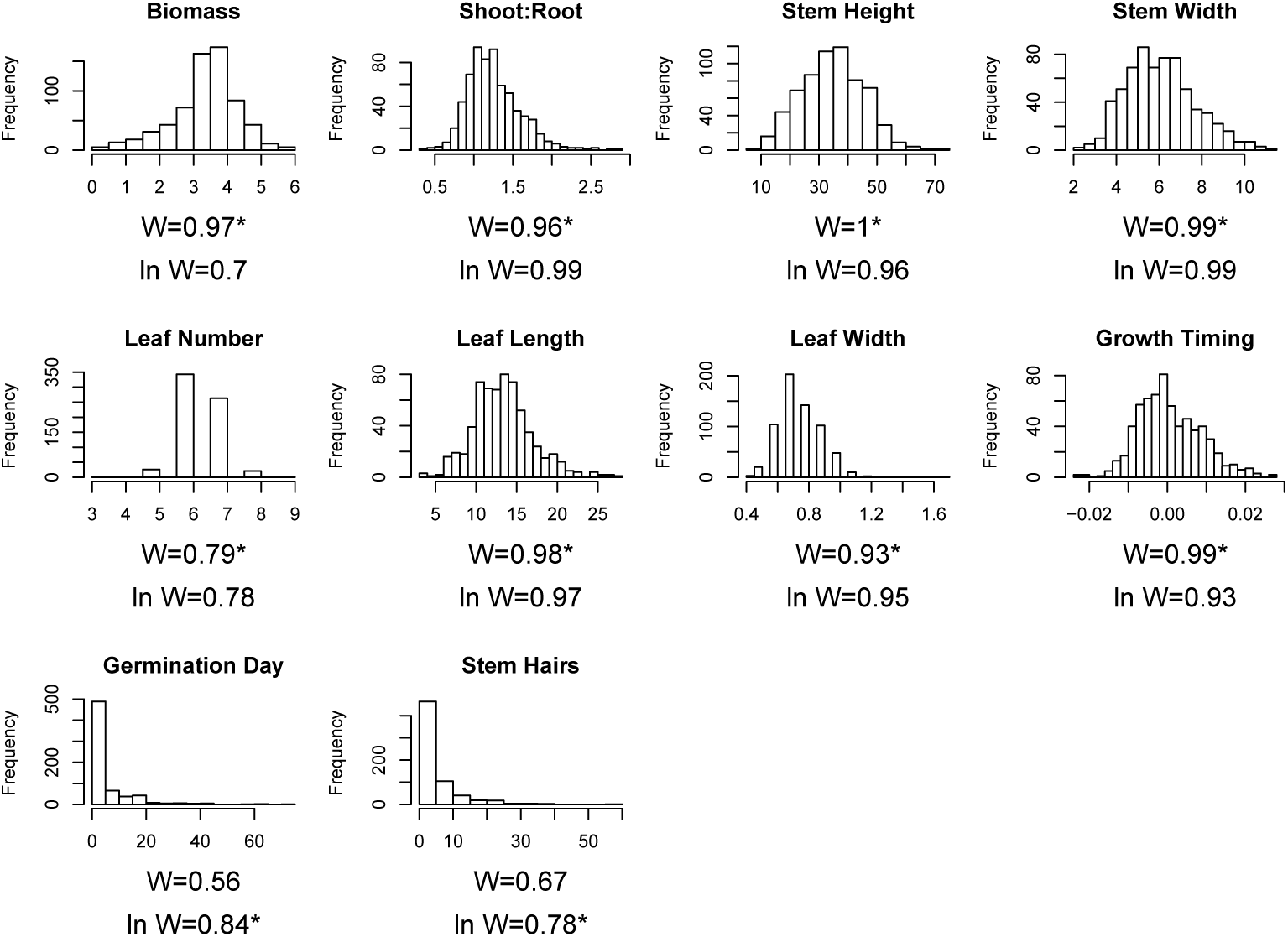
Histogram of measured data for traits. W-statistics from Shapiro tests are included, as well for Shapiro tests for the natural log of the data (plus 1 for Stem Hairs to retain observations with 0 hairs as datapoints), and an asterisk marks which distribution (raw or natural log) was used in analyses.

**Figure S5:**
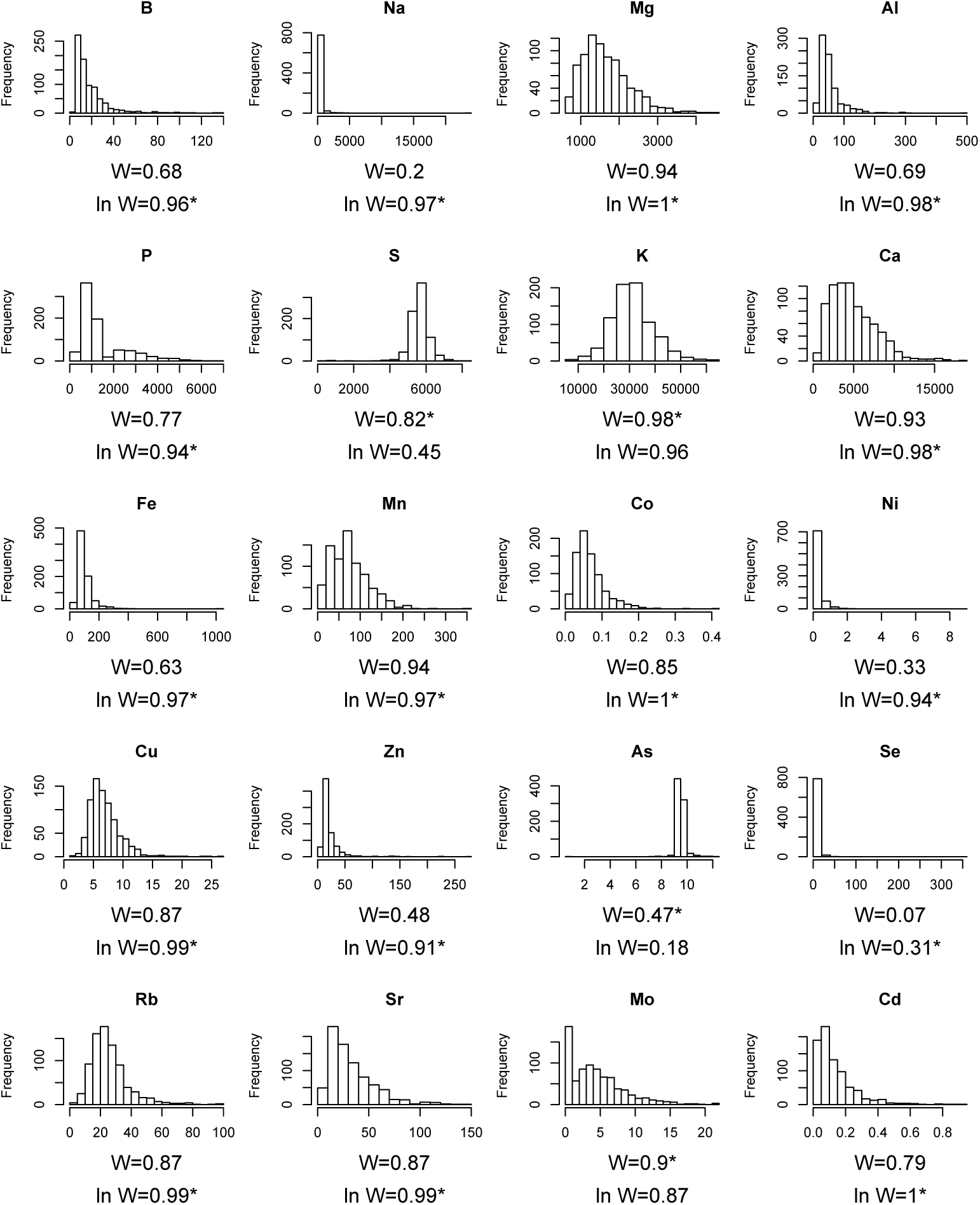
Histogram of measured data for tissue element concentrations (by weight). W-statistics from Shapiro tests are included, as well for Shapiro tests for the natural log of the data, and an asterisk marks which distribution (raw or natural log) was used in analyses.

**Figure S6:**
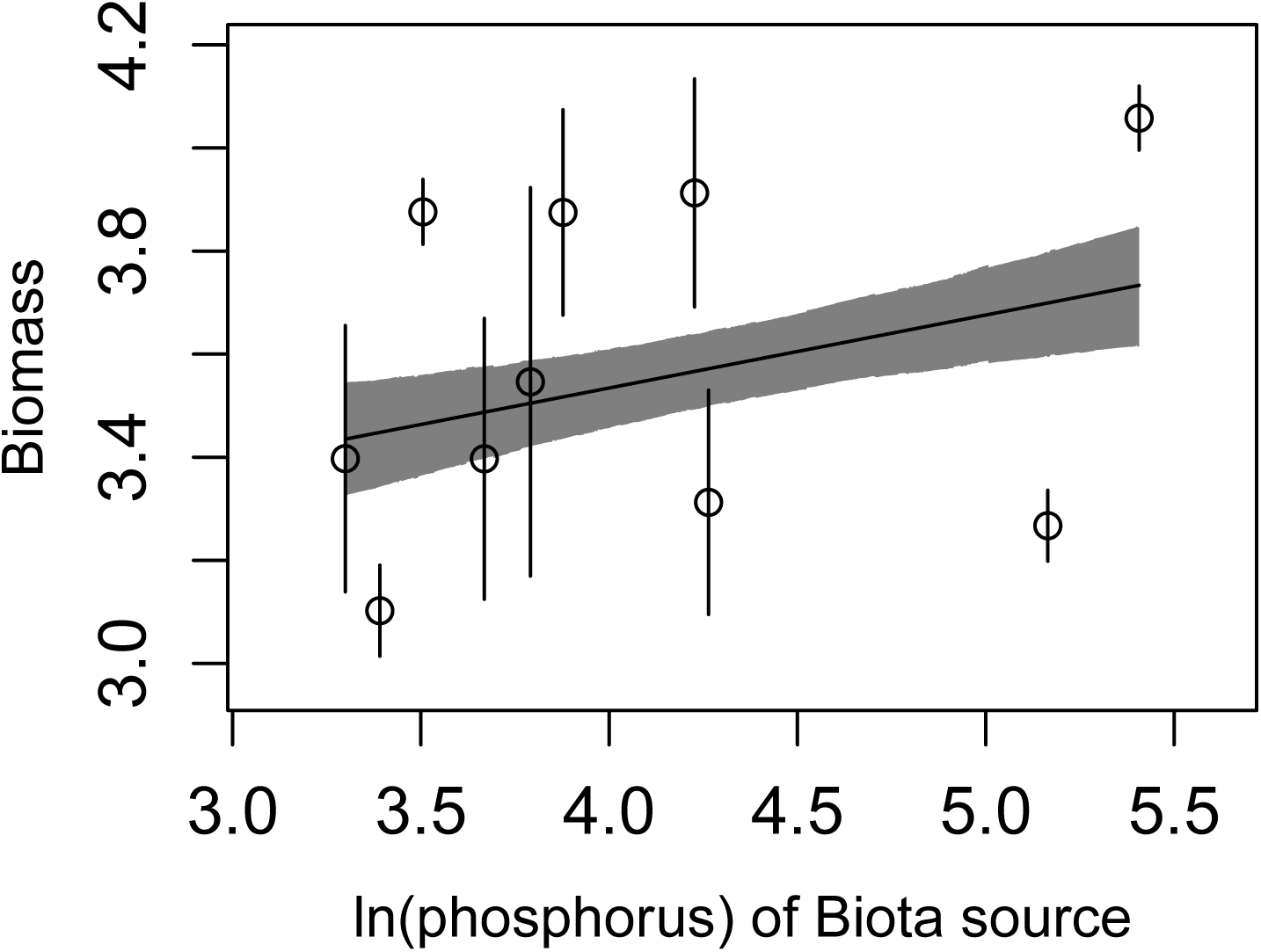
Best model of those including phosphorus at the source site as the explanatory environmental variable. Note this model fits worse than the model using mean annual temperature, but this figure is included for comparison. Only *β_E_B__* is significant.

**Figure S7:**
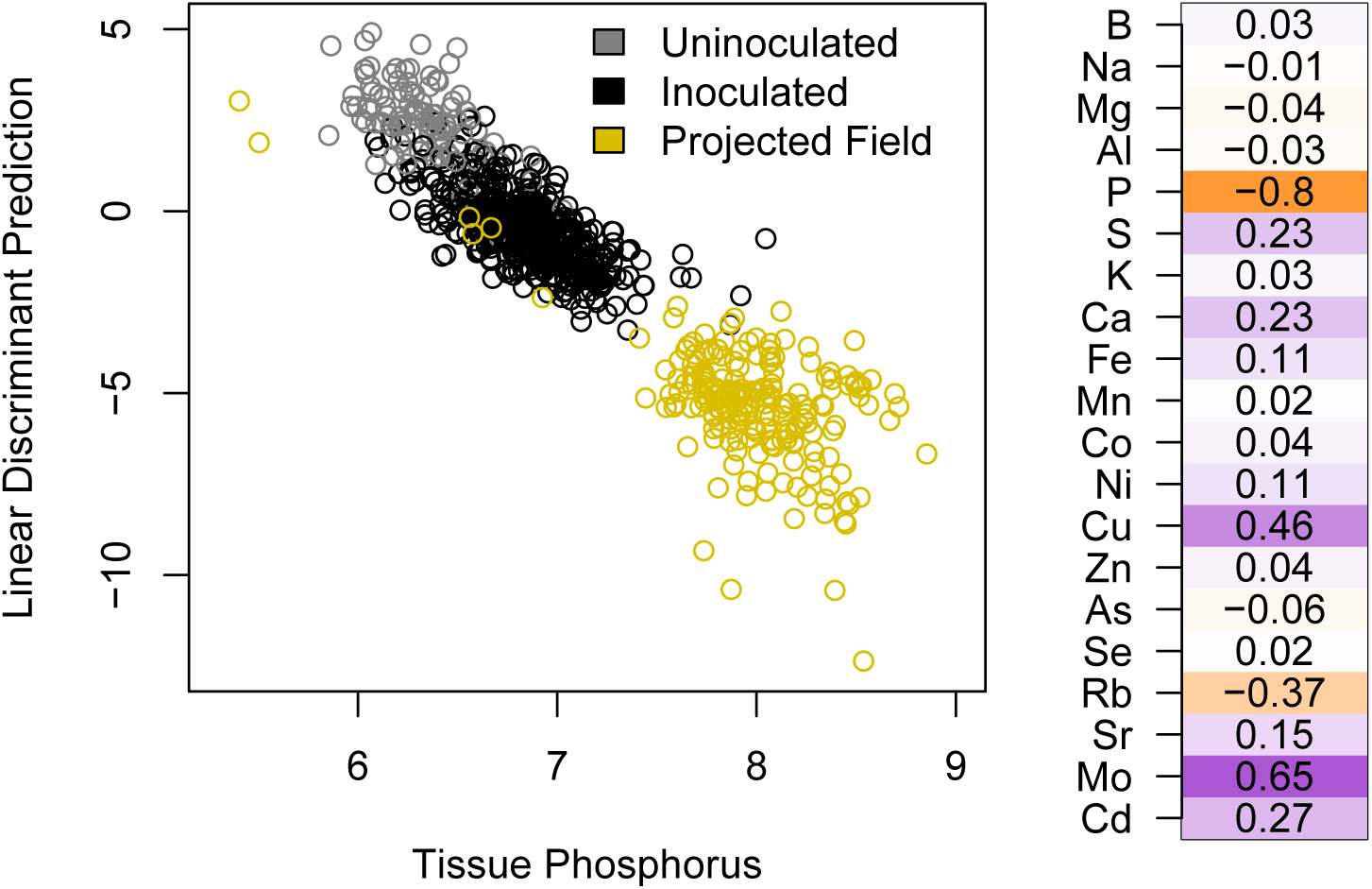
Left, linear discriminant analysis of elemental profiles between inoculated (black) and uninoculated (grey) plotted against tissue phosphorus (logged values). Projections for field plants in yellow. Right, correlations of individual element concentrations with the resulting LDA prediction scores in orange (negative) or purple (positive), with stronger colors indicating stronger *ρ*.

**Figure S8:**
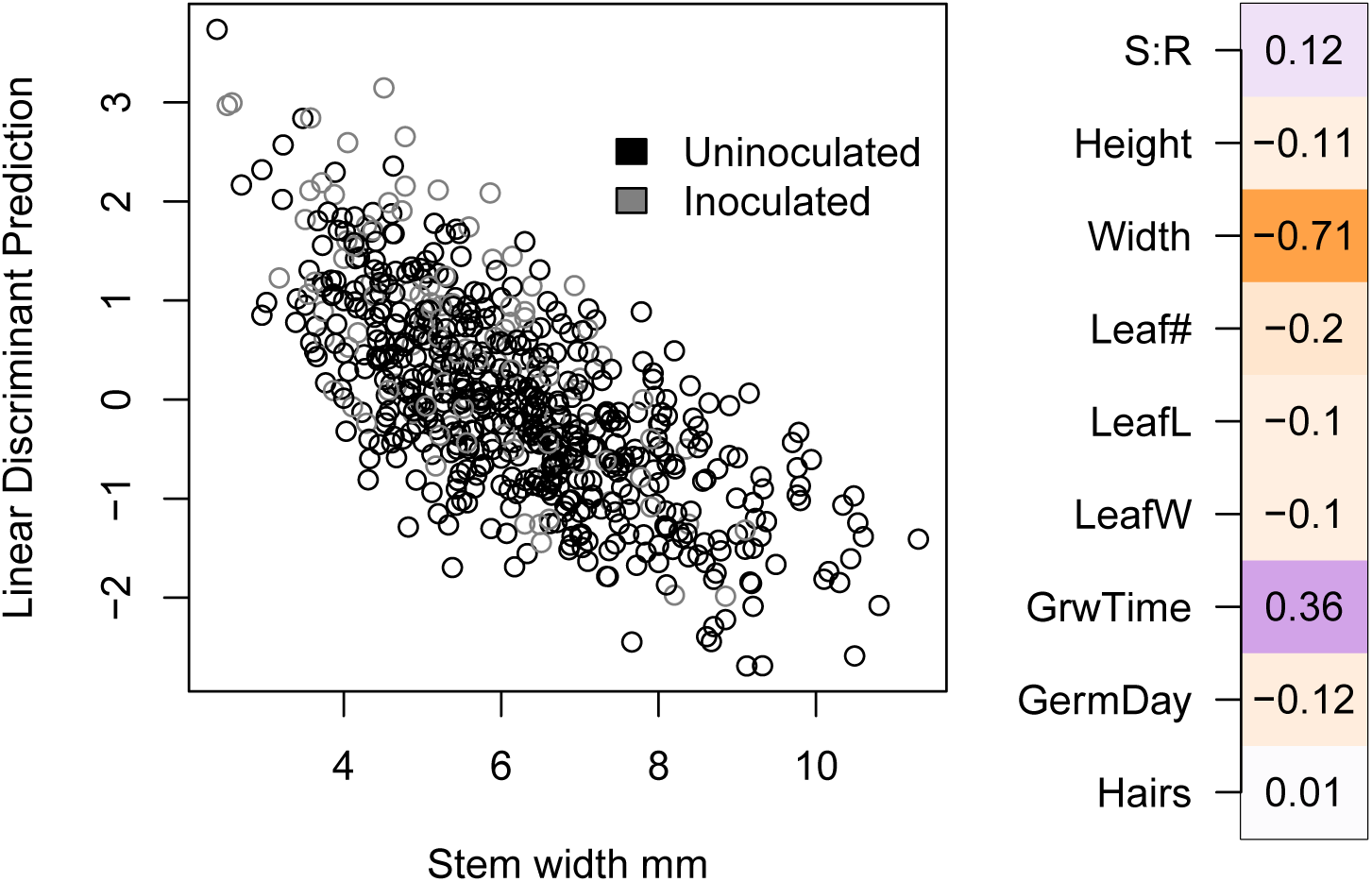
Left, linear discriminant analysis of traits between inoculated (black) and uninoculated (grey) plants relies primarily on stem width. Right, correlations of individual traits with the resulting LDA prediction scores in orange (negative) or purple (positive), with stronger colors matching strength of *ρ*.

**Figure S9:**
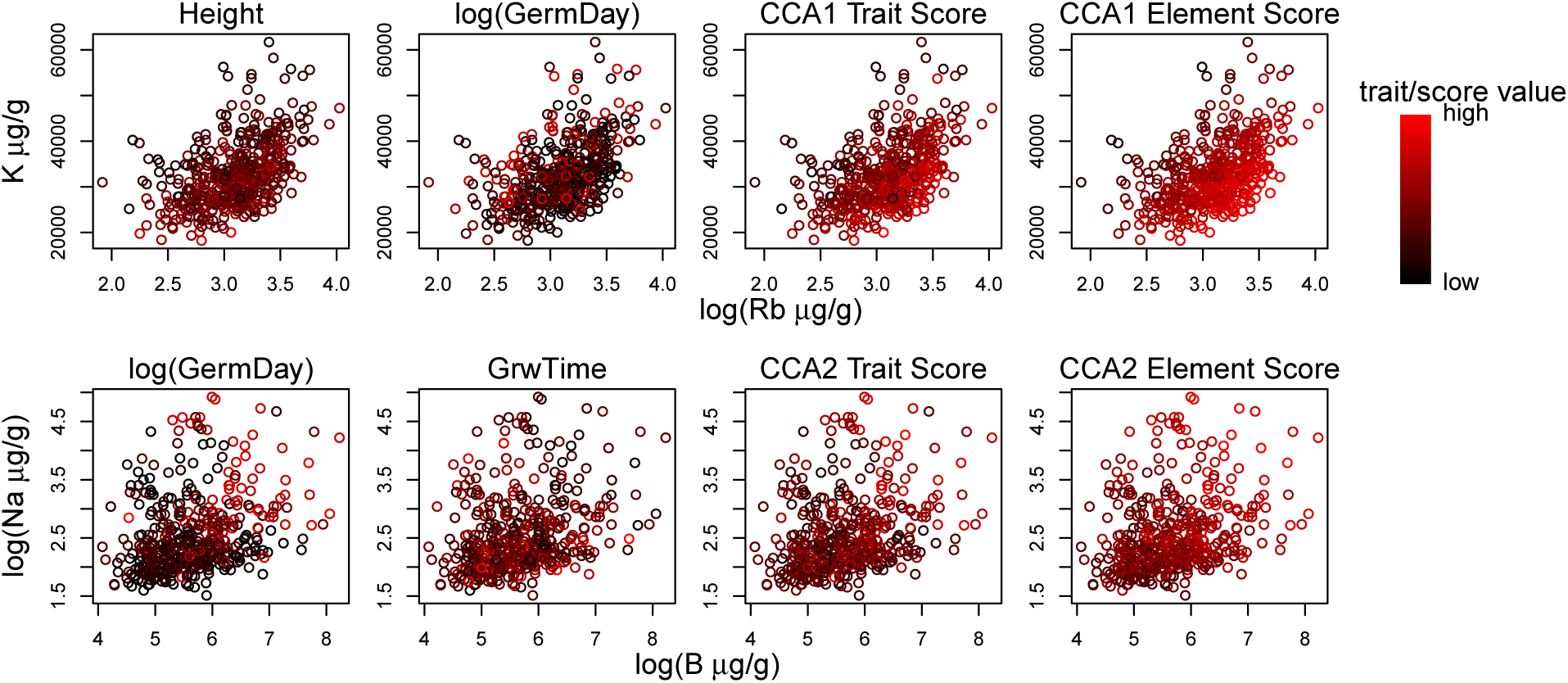
Visualizng a subset of the multivariate relationships identified by the CCA. CCA axes identify correlations in highly multivariate space; meaning, they often include shifts in the relative concentrations of elements to each other, or relative values of traits to each other. Plots in the upper row some of the relationships identified in the first CCA axis: rubidium and plant height load strongly on this axis in the positive direction, and most other elements (potassium here, for example), as well as the log of days until germination load in the negative direction. Plants that were relatively higher in rubidium for a given concentration of potassium (points shifted towards the upper left corners of plots) were taller, germinate earlier, and had higher scores on the CCA axis. Plots in the lower row show examples of the multivariate relationships identified by the second CCA axis: boron, sodium, and the log of germination day load strongly in the positive direction on this axis, while the the timing of vegetative growth loads negatively. Plants that had higher levels of both boron and sodium in tissues (those in the top right corners only) germinated later but grew fastest right after germinating. See Figure 2 for complete multivariate CCA axis loadings. Abbreviations: log(GermDay), natural log of days until germination; GrwTime, timing of vegetative growth; standard elemental abbreviations. The natural log of elemental concentrations is shown here when logged data were used in the analyses (Figure S5).

**Figure S10:**
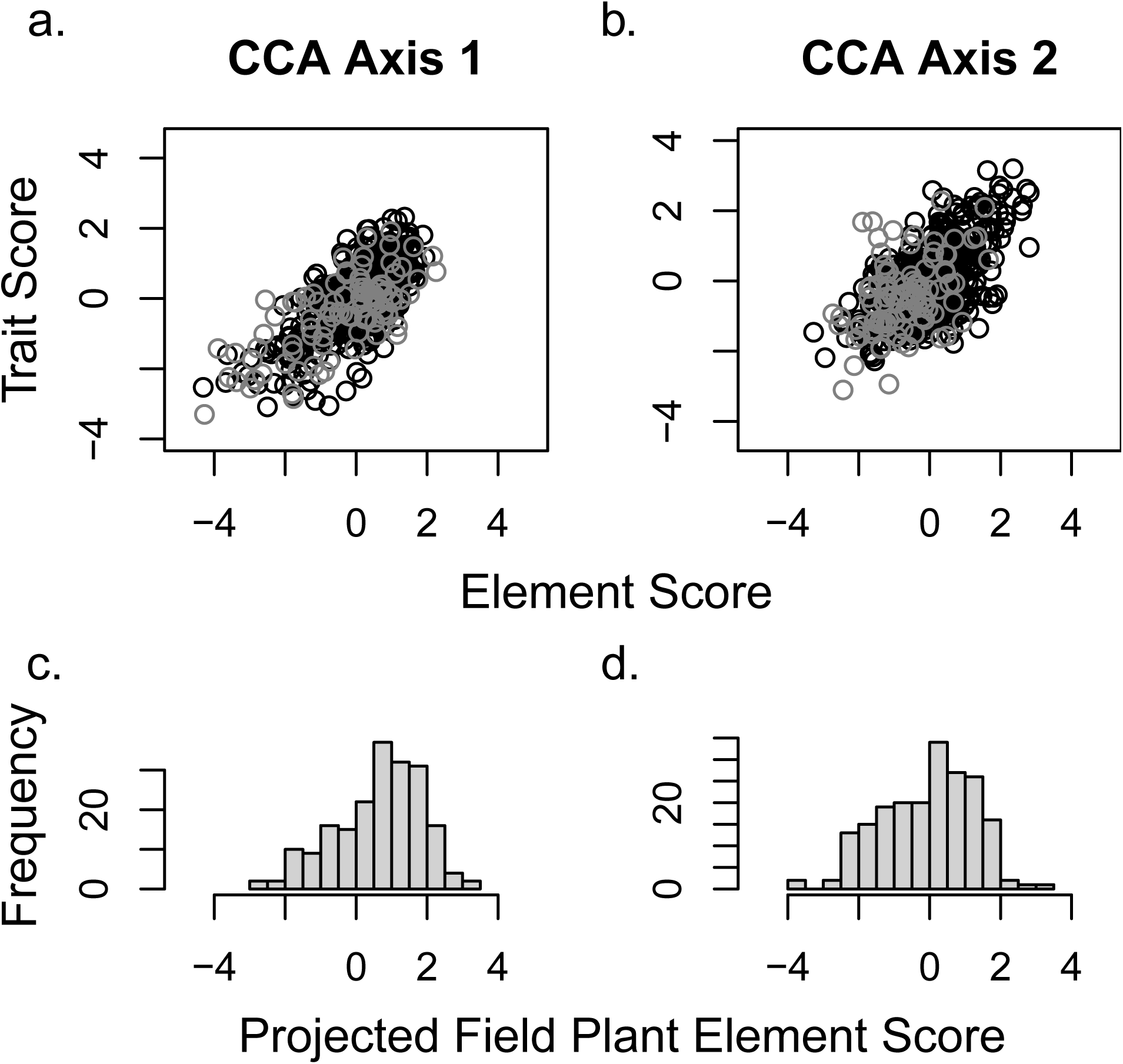
Projections of field and uninoculated greenhouse plants onto the CCA axes. CCA plots of both x and y variables for axis 1 (a) and 2 (b), see also 2, showing inoculated greenhouse plants in black, and uninoculated greenhouse plants in grey. Histograms of x-axis projections of elements measured in field plants (full trait data were not available for field plants) for first (c) and second (d) CCA axes.

**Figure S11:**
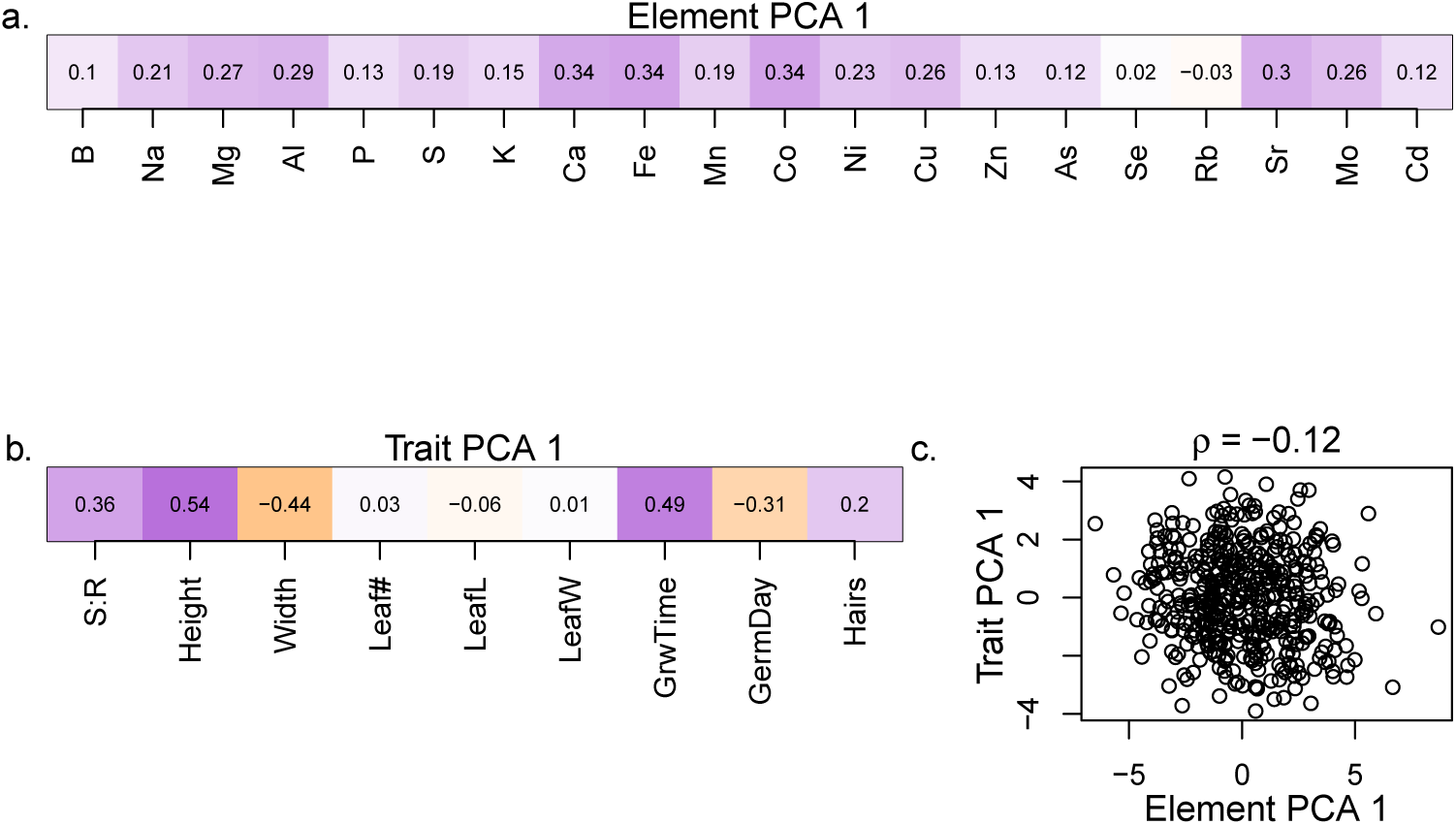
Loadings of elements onto the first axis of the PCA of element profile data alone (a). Loadings of traits onto the first axis of the PCA of trait data alone (b). In (c), scores for plants on the first axis of each respective PCA are plotted against each other, showing no strong relationship. Abbreviations of traits and elements are as elsewhere.

**Figure S12:**
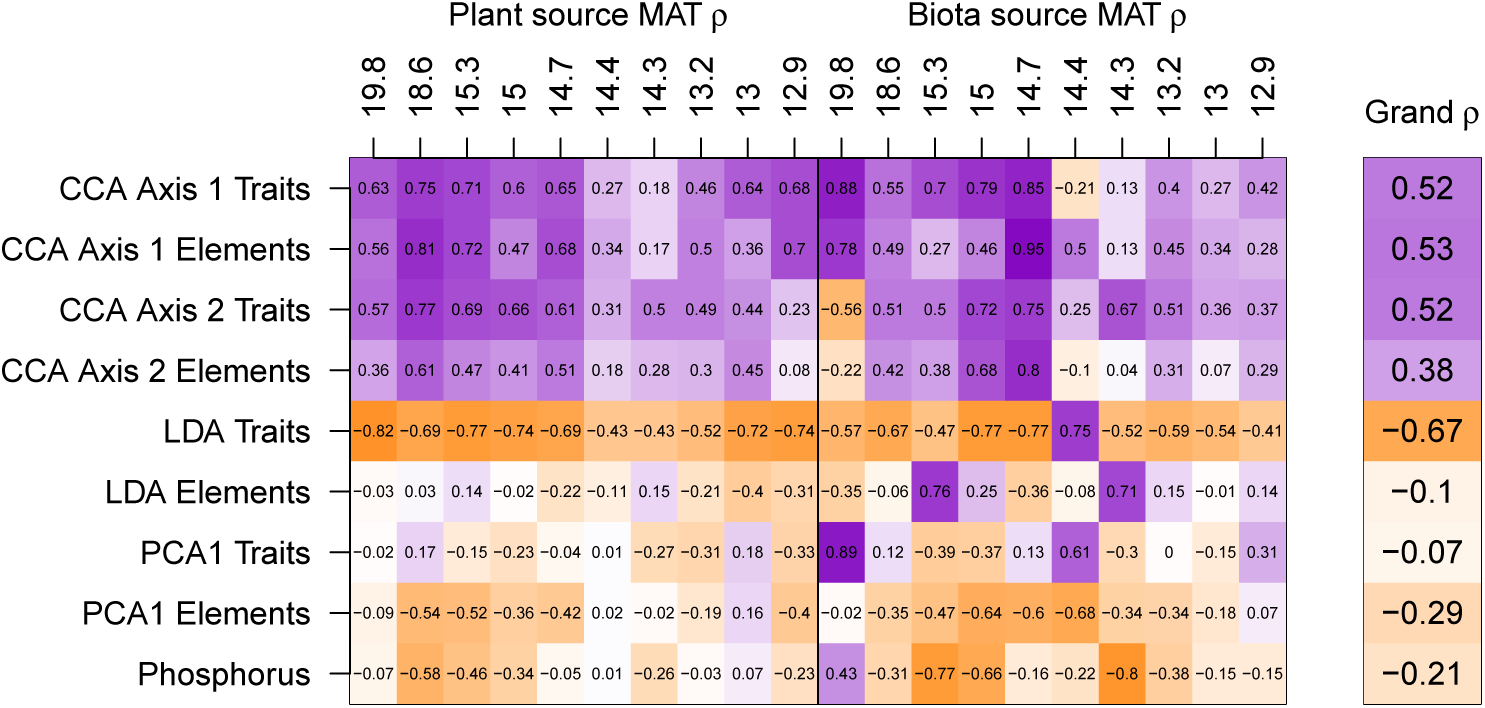
Correlation of multivariate axes with biomass three different ways: within plant source mean annual temperature (MAT, but across families, replicates and biota source MAT), within biota source MAT (but across families, replicates and plant source MAT), and across all data (Grand *ρ*). Note that correlations within biota source MAT are based on sympatric combinations only (plants from the same site) for most sites (MAT 15, 14.7, 14.4, 14.3, 13.2, and 12.9*^◦^*C). Colors are purple for positive correlations and orange for negative correlations, with color strength reflecting strength of correlation.

**Figure S13:**
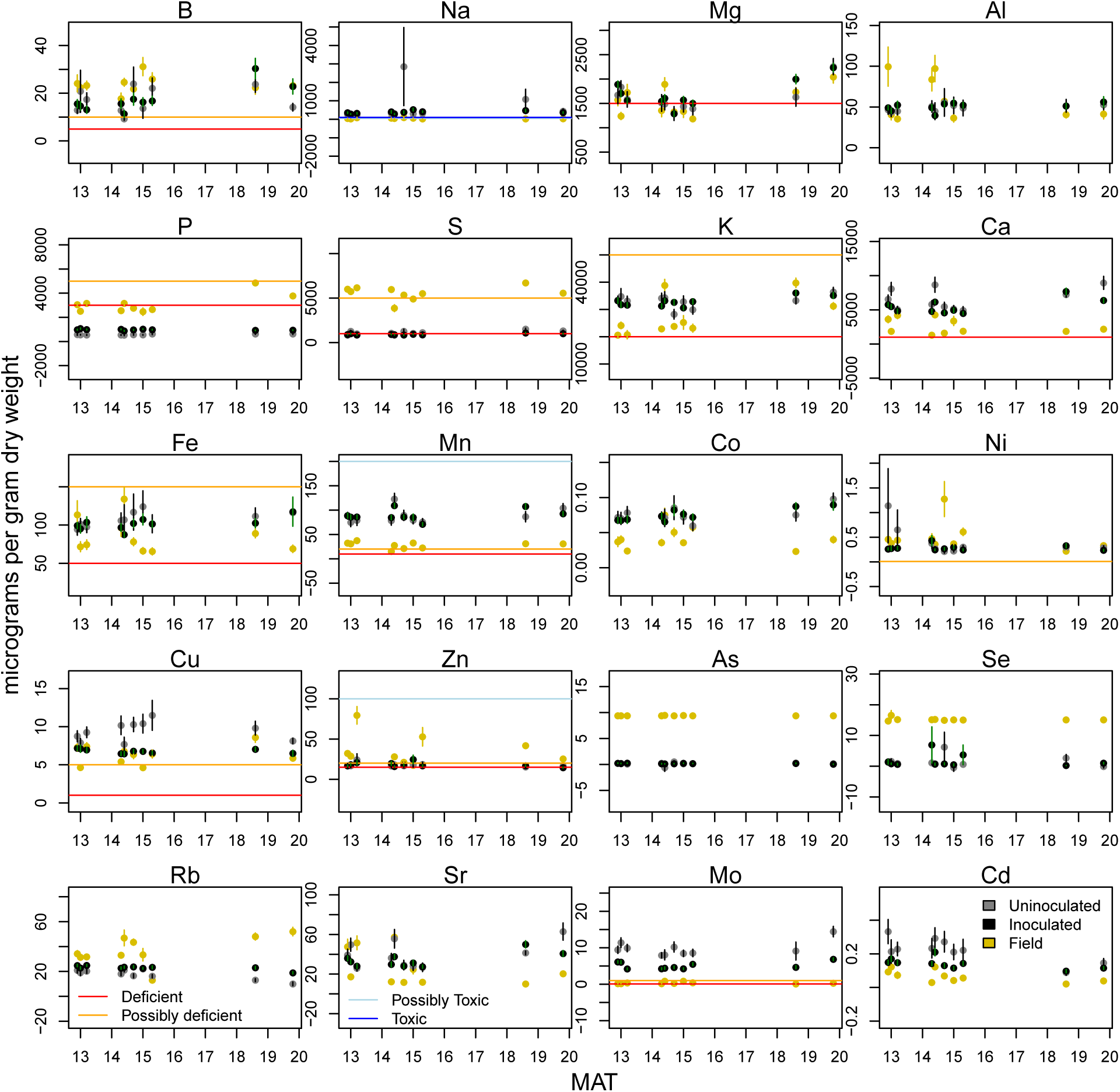
Average measured tissue element concentrations (*µ*g g*^−^*^1^ using dry weight) for field (yellow), greenhouse inoculated (black) and uninoculated (grey) plants plotted against mean annual temperature of the field site. Vertical bars indicate one standard error of the mean. Horizontal lines indicate a value at which that element is very likely (red) or possibly (orange) limiting growth due to deficiency, or where that element is very likely (dark blue) or possibly (light blue) limiting growth due to toxicity (values from maize, grasses or plants broadly as available in Marschner, 2011). In many cases, one or more thresholds are far from actual tissue concentrations and are not visible. Seven elements (Al, Co, Se, Rb, Sr, and Cd) have no visible thresholds because they have no, or uncertain, beneficial concentrations and are not near any toxicity threshold.

**Table S1:** Sampling site abiotic characteristics. Climate and elevation data downloaded from BioClim (Hijmans et al., 2005) extracted with raster in R (Hijmans, 2015). MAT, mean annual temperature in *^◦^*C; TAP, total annual precipitation in mm; SWC (%), soil water holding capacity. Soil testing service was purchased from INIFAP in Celaya, Gto, methods as provided. Inorganic N was KCl extraction with MgO distilation, P was quantified with the Bray method. K, Ca, Mg, and Na were extracted in ammonium acetate 1N at pH7, while Fe, Mn, Cu, and Zn were extracted with diethylenetriaminepentaacetic acid. Both metal groups were quantified with atomic absorption or inductively coupled plasma. Elevation is in meters above sea level Soil elements are in micrograms per gram of dry weight (equivalently, ppm). A * indicates sites used as rhizosphere inocula across all populations of teosinte. A indicates measurements or extracted variables also reported in O’Brien et al. (2019). Growing season for teosinte is in the warmer, wetter, portion of the year. Greenhouse average temperature from first possible germination day to last harvest day was 23.8 ◦C, average night temperature for the same period was 19.4 ◦C, and relative humidity average was 61.7.

|  | Calimaya<br>Upper | Toluca* | Calimaya<br>Lower | San<br>Matías<br>Cuijingo | San<br>Francisco<br>Pedregal | Tenango<br>Del<br>Aire | San<br>Mateo<br>Tezoquipan | Texcoco* | Malinalco* | Tepoztlán* |
| --- | --- | --- | --- | --- | --- | --- | --- | --- | --- | --- |
| Longitude | -99.633 | -99.722 | -99.616 | -98.843 | -99.127 | -98.863 | -98.809 | -98.922 | -99.501 | -99.070 |
| Latitude | 19.161 | 19.260 | 19.151 | 19.077 | 19.212 | 19.146 | 19.212 | 19.505 | 18.954 | 18.976 |
| MAT† | 12.9 | 13.0 | 13.2 | 14.3 | 14.4 | 14.7 | 15.0 | 15.3 | 18.6 | 19.8 |
| TAP† | 857 | 836 | 828 | 926 | 935 | 817 | 730 | 585 | 928 | 966 |
| Elevation† | 2792 | 2776 | 2698 | 2491.5 | 2507 | 2408 | 2353 | 2253 | 1881 | 1665 |
| P (Bray) | 71.1 | 29.7 | 39.2 | 27.1 | 44.3 | 48.3 | 68.5 | 175 | 223 | 33.3 |
| Inorganic N | 16.2 | 17.6 | 14.1 | 12.7 | 13.4 | 15.5 | 12.0 | 13.4 | 16.9 | 12.0 |
| SWC† | 23.3 | 33.0 | 21.8 | 33.0 | 33.8 | 27.8 | 30.8 | 40.5 | 55.5 | 30.0 |
| K | 142 | 143 | 96.3 | 261 | 498 | 189 | 315 | 1055 | 827 | 428 |
| Ca | 749 | 1181 | 354 | 1034 | 966 | 757 | 1575 | 2710 | 4076 | 915 |
| Mg | 26.2 | 286 | 55.5 | 170 | 167 | 240 | 227 | 660 | 527 | 290 |
| Na | 25.9 | 34.3 | 26.4 | 10.3 | 27.5 | 10.7 | 14.8 | 419 | 31.5 | 28.2 |
| Fe | 31.8 | 176 | 25.9 | 64.8 | 34.5 | 56.0 | 53.8 | 29.6 | 81.8 | 58.6 |
| Zn | 0.67 | 3.25 | 0.39 | 1.24 | 2.63 | 1.81 | 7.17 | 10.5 | 48.7 | 1.94 |
| Mn | 4.63 | 75.0 | 2.23 | 5.95 | 2.51 | 5.94 | 6.02 | 6.28 | 20.5 | 15.7 |
| Cu | 0.38 | 1.49 | 0.23 | 0.97 | 1.07 | 0.88 | 1.4 | 2.12 | 0.96 | 1 |

**Table S2:** Sampling site biotic characteristics. Coarse phenology was approximate qualitative percentages at two metrics scored by visually estimating the population at the time of seed collection. Plants that remained at least partly green were “Alive” (others were fully senesced). Plants with undeveloped fruit (even if also fully dead) were “Immature Fruit”(%Immat Fruit), while other plants were either empty of fruit or had mature fruits (%Mat Fruit). Spore counts are per gram of soil used to mix inocula. A * indicates sites used as rhizosphere inocula across all populations of teosinte. Populations are sorted by increasing MAT, see Table S1

|  | Calimaya<br>Upper | Toluca* | Calimaya<br>Lower | San<br>Matías<br>Cuijingo | San<br>Francisco<br>Pedregal | Tenango<br>Del<br>Aire | San<br>Mateo<br>Tezoquipan | Texcoco* | Malinalco* | Tepoztlán* |
| --- | --- | --- | --- | --- | --- | --- | --- | --- | --- | --- |
| %Alive | 8 | 5 | 2 | 5 | 5 | 2 | 10 | 0 | 0 | 0 |
| %Immat Fruit | 45 | 35 | 35 | 15 | 20 | 10 | 10 | 85 | 10 | 5 |
| %Mat Fruit | 50 | 60 | 60 | 80 | 75 | 80 | 80 | 10 | 75 | 75 |
| Spore count | 1233 | 159 | 532 | 915 | 762 | 432 | 223 | 174 | 567 | 419 |

**Table S3:** All element concentrations are in micrograms per gram dry weight, but were logged where this improved normality (based on the W statistic of a Shapiro test in R, those not logged marked with ; see main text, Figures S4, S5. Intercepts are significantly different from 0, unless indicated with “n.s.”. For slopes pMCMC of interest are indicated with: *** is *<* 0.001, ** *<* 0.01, * is *<* 0.05, . is *<* 0.1. Models were fit with MCMCglmm (Hadfield, 2010), with 13,000 iterations, 3,000 burn-in, and thining by 10. Note pMCMC values are not multiple-test corrected.

| Trait or element | Live intercept | Live Slope | Sibling Intercept | Sibling Slope |
| --- | --- | --- | --- | --- |
| Biomass | 2.58 | 0.91*** | 3.48 | -0.56*** |
| Shoot:Root | 1.28 | -0.028 | 1.28 | 0.008 |
| Height cm | 34.20 | 0.55 | 28.22 | 3.02*** |
| Stem width mm | 5.48 | 0.75*** | 7.26 | -0.79*** |
| Leaf Number | 6.34 | 0.097 | 6.54 | -0.002 |
| Leaf Length cm | 13.09 | 0.28 | 10.67 | 0.49*** |
| Leaf Width cm | 0.75 | 0.013 | 0.59 | 0.027* |
| Growth Acceleration | 0.0025 | -0.0022** | 0.0012 n.s. | 0.0002 |
| Germination Day† | 1.37 | 0.10 | 1.15 | 0.035*** |
| Hairs per cm<sup>2</sup>† | 0.97 | 0.013 | 0.80 | 0.045*** |
| B | 2.60 | -0.046 | 2.03 | 0.18** |
| Na | 5.62 | 0.011 | 5.48 | 0.029 |
| Mg | 7.33 | 0.031 | 5.30 | 0.28*** |
| Al | 3.76 | 0.032 | 3.71 | 0.020 |
| P | 6.32 | 0.53*** | 7.08 | -0.037 |
| S† | 5778 | -172*** | 4683 | 0.16*** |
| K† | 33086 | -0383 | 29500 | 0.097. |
| Ca | 8.71 | -0.21*** | 6.67 | 0.21*** |
| Fe | 4.61 | -0.079* | 4.50 | 0.0065 |
| Mn | 4.42 | -0.016 | 3.37 | 0.23*** |
| Co | -2.67 | -0.042 | -2.59 | 0.059 |
| Ni | -1.34 | -0.13* | -1.49 | 0.019 |
| Cu | 2.17 | -0.31*** | 1.63 | 0.10* |
| Zn | 2.80 | -0.039 | 3.01 | -0.087 |
| As† | 9.48 | 0.063 | 9.27 | 0.03 |
| Se | 2.77 | -0.0063 | 2.19 | 0.20*** |
| Rb | 2.81 | 0.27*** | 2.61 | 0.17*** |
| Sr | 3.53 | -0.17** | 2.75 | 0.17*** |
| Mo† | 9.78 | -4.72*** | 3.85 | 0.11*** |
| Cd | -1.78 | -0.41*** | -1.71 | 0.27*** |

**Table S4:** Biota and plant source effects on plant values for select other response values including: the top trait on each of the first two CCA axes between trait and element matrices (height and germination), stem width (since it was measured in the field and somewhat strongly correlated to CCA2), the first axis of each PCA for element and trait values separately (agnostic approach for trait-ion linkage), and LDA axes. Abbreviations: elmt is element; trt is trait; Germ is the natural log of the germination day, and StemW is Stem width. Intercepts, representing values for 0 MAT, are not meaningful. –: not included in best model **: pMCMC *<* 0.01,*: pMCMC *<* 0.05.

|  |  | CC1 elmt | CC2 elmt | Height | Germ | StemW | LDA elmt | LDA trt | PCA1 trt | PCA1 elmt |
| --- | --- | --- | --- | --- | --- | --- | --- | --- | --- | --- |
| Intercept | $a$ | -0.33 | -2.75** | 75.7** | -0.91** | -2.24** | 0.18 | 4.17** | 8.22** | -0.62 |
| Biota source Env. | $\beta_B$ | 0.18** | 0.068** | — | 0.0012 | 0.020** | -0.027** | -0.019** | -0.0097** | -0.012** |
| Plant source Env. | $\beta_P$ | -0.17** | 0.11** | -0.26** | 0.017** | 0.034** | 0.0099. | -0.0075** | -0.044** | 0.017** |
| Sympatry | $\beta_S$ | 1.63* | — | — | — | 2.28* | — | -2.16** | — | — |
| Sympatry × Env. | $\beta_{E \times S}$ | -0.097* | — | — | — | -0.014* | — | 0.013** | — | — |
| Best Env. |  | MAT | MAT | MAT | MAT | MAT | SWC | MAT | MAT | MAT |

**Table S5:** Best models fitted to selected further response variables for plants in the field: projections of the field elemental profiles onto the LDA axis and first PCA axis for elemental profiles from the greenhouse data. For each response variable, we report the model coefficients of the best model in rows. Significance of intercepts is not meaningful, representing values for 0 ◦C. –: not included in best model, **: pMCMC *<* 0.01, *: pMCMC *<* 0.05, .: pMCMC *<* 0.1.

| | Intercept | $\beta_{MAT}$ | $\beta_C$ | $\beta_{MAT \times C}$ |
| --- | --- | --- | --- | --- |
| **LDA** | 2.29* | -0.014** | -2.05.* | – |
| **PCA** | -2.60** | – | – | – |

## Footnotes

1 Significance of intercepts are not meaningful (representing 0 ◦C MAT). –: not included in best model **: pMCMC < 0.01,*: pMCMC < 0.05, . : pMCMC < 0.1.

2 *β*_MAT_ is substituted for *β*_E_, as MAT was the only variable tested for field data (see Methods). Significance of intercepts is not meaningful, representing 0◦C. –: not in best model, **: pMCMC < 0.01, *: pMCMC < 0.05, .: pMCMC < 0.1.

## Notes

### Competing Interest Statement

The authors have declared no competing interest.

